# LRBA balances antigen presentation and T-cell responses *via* autophagy by binding to PIK3R4 and FYCO1

**DOI:** 10.1101/2022.10.17.512524

**Authors:** Elena Sindram, Marie-Celine Deau, Laura-Anne Ligeon, Pablo Sanchez-Martin, Sigrun Nestel, Sophie Jung, Stefanie Ruf, Pankaj Mishra, Michele Proietti, Stefan Günther, Kathrin Thedieck, Eleni Roussa, Angelika Rambold, Christian Münz, Claudine Kraft, Bodo Grimbacher, Laura Gámez-Díaz

## Abstract

Reduced autophagy is associated with the aberrant humoral response observed in lipopolysaccharide-responsive beige-like anchor protein (LRBA) deficiency; however, the exact molecular mechanism and its impact on T-cell responses remain unknown. We identified two novel LRBA interactors, phosphoinositide 3-kinase regulatory subunit 4 (PIK3R4) and FYVE And Coiled-Coil Domain Autophagy Adaptor 1 (FYCO1). Both proteins play essential roles in different stages of autophagy. PIK3R4 facilitates the production of phosphatidylinositol-3 phosphate (PI(3)P) required for autophagosome formation and autophagosome-lysosome fusion, whereas FYCO1 allows autophagosome movement. LRBA-KO cells showed an impaired PI(3)P production, a delayed autophagosome-lysosome fusion, an accumulation of enlarged autophagosomes, and an atypical lysosomal positioning. These abnormalities led to decreased cargo material degradation and prolonged antigen presentation to T-cells via autophagy, resulting in increased production of proinflammatory cytokines, as autophagy is a major intracellular degradation system for major histocompatibility class II complex (MHCII) loading. Aberrant autophagosome formation, cargo degradation and antigen presentation were rescued by ectopic expression of WT-LRBA. In summary, we identified a novel function of LRBA that is crucial for T-cell-driven response through the interaction with two proteins of the autophagy machinery. These observations may contribute to the exacerbated T-cell dysregulation observed in LRBA-deficient patients.

## Introduction

LRBA deficiency is an inborn error of immunity (IEI) caused by deleterious biallelic mutations in *LRBA* (*1*). LRBA-deficient patients display a broad spectrum of manifestations ranging from immunodeficiency to T-cell driven immune dysregulation (*2–5*). The latter is associated with reduced cytotoxic T lymphocyte-associated protein 4 (CTLA-4) expression on regulatory T cells (Tregs) due to defective CTLA-4 recycling orchestrated by LRBA (*6*). Therefore, an overlap between the clinical pictures of LRBA deficiency and CTLA-4 insufficiency is frequently observed (*7, 8*). However, abnormal CTLA-4 trafficking does not explain the increased severity, poor survival and earlier onset of symptoms observed in LRBA deficiency compared to CTLA-4 insufficiency patients. We have previously reported that B lymphocytes from LRBA-deficient patients exhibit defective autophagy leading to increased apoptosis of plasmablasts, explaining the poor humoral immune response (*1*). Autophagy is a conserved intracellular degradation process essential for cell survival, by which cytoplasmic material, including aberrant proteins or intracellular pathogens, are engulfed in double-membrane vesicles called autophagosomes that subsequently fuse with lysosomes for cargo degradation (*9*). Although autophagy is a constitutively active housekeeping mechanism that promotes cell survival, it is also essential in immune cells for modulating chemokines release upon antigen presentation restricted to MHC-II (*10–12*).

To date, the molecular mechanism of LRBA in autophagy and the impact of LRBA loss in autophagy-dependent immune functions remain, however, unknown. In this study, we identified a role of LRBA at early and late stages of autophagy by physically interacting with PIK3R4 and FYCO1. PIK3R4 is the regulatory subunit of the phosphatidylinositol 3-kinase (PI3K-III) complex that positively regulates autophagy through the formation of phosphatidylinositol-3-phosphate (PI(3)P). In turn, PI(3)P allows the recruitment of double FYVE domain-containing protein-1 (DFCP-1) and WD repeat domain phosphoinositide-interacting protein-2 (WIPI2) to initiate the formation of autophagosomes (*13*). FYCO1 functions as an adapter protein linking autophagosomes to microtubule plus-end-directed molecular motors, allowing autophagosome movement after binding PI(3)P (*14, 15*). We demonstrated that LRBA regulates PI(3)P production, recruitment of DFCP-1 and WIPI2, size and movement of autophagosomes, autophagosome/lysosome fusion, and cargo degradation *via* autophagy. Additionally, we observed that LRBA is essential for the regulation of MHC-II-restricted antigen presentation, thereby regulating T-cell proinflammatory cytokines release. Finally, the reconstitution of LRBA in LRBA-KO cells re-established the normal autophagy flux and antigen presentation, validating the specific role of LRBA in autophagy. Taken together, our data indicate that LRBA forms different protein complexes serving at different stages of autophagy, and that loss of LRBA impacts the targeting of cytosolic antigens for cargo degradation *via* autophagy, enhancing antigen presentation. The latter may contribute to the exacerbated T-cell immune dysregulation observed in LRBA-deficient patients.

## Results

### LRBA interacts with PIK3R4

Using computational predictions based on STRING and FuncBase databases (*16, 17*), we identified 28 potential LRBA interactors, which were found to be enriched in autophagy and vesicle-trafficking events according to Gene Ontology (GO) –term enrichment analysis (Fig.1A, Table 1). Only PIK3R4 was predicted to interact with LRBA by both databases. The interaction of LRBA and PIK3R4 was validated at endogenous levels in HEK293T cells by co-immunoprecipitation (co-IP) (fig. S1A) and in lymphoblastoid B cell lines (LCL) using a proximity ligation assay (PLA) (Fig. 1, B and C). In both assays, LRBA-KO HEK293T cells or LCL cells from LRBA-deficient patients were used as controls as no LRBA signal was observed. The LRBA: PIK3R4 interaction was also validated in an overexpression system by co-IP in HEK293T cells previously co-transfected with myc-tagged LRBA and his-tagged PIK3R4 (Fig. 1D). Co-IP experiments using seven different flag-tagged LRBA protein domains (F1-F7) and myc-tagged PIK3R4 identified that the LRBA WD40 domain interacts with PIK3R4 (Fig.1, E and F). We prepared a structural model of LRBA and the PIK3R4/PIK3-III complex by homology modelling. The non-complexed homology model of the PIK3R4 WD40 domain exhibited one single helix on the surface that was defined as interacting constraints for the docking protocol (Fig. 1G). The resulting models of the HADDOCK web server indicated one top-ranked cluster with a high negative Z-score value (-2.0). In the best model, the exposed ɑ-helix of PIK3R4 interacts with the upper surface of the β-propeller via hydrophobic and charged interactions (Fig. 1H). Together with the PI3K catalytic subunit type 3 (better known as VPS34) and Beclin-1, PIK3R4 forms the core of the PI3K-III complex. Throughout the different stages of autophagy, the PI3K-III complex recruits different adaptor proteins, including AuTophaGy Related protein 14L (ATG14L) and UV radiation resistance-associated gene protein (UVRAG) (*18, 19*). Following PLA and co-IP assays to evaluate potential interactions of LRBA with other members of the PI3K-III complex, we observed an interaction of LRBA with UVRAG (fig. S1B) but not with VPS34, Beclin-1, or ATG14L (fig. S1, C-E). Moreover, the absence of PLA signals supported no physical proximity of LRBA with PI3K-delta, another member of the PI3K family that is essential for immune responses (fig. S1F). Altogether, these results demonstrate that LRBA interacts specifically with PIK3R4 from the PI3K-III complex through the WD40 domain, possibly influencing UVRAG-containing PI3K-III complexes that are involved in autophagosome maturation (*20*).

**Fig. 1.**
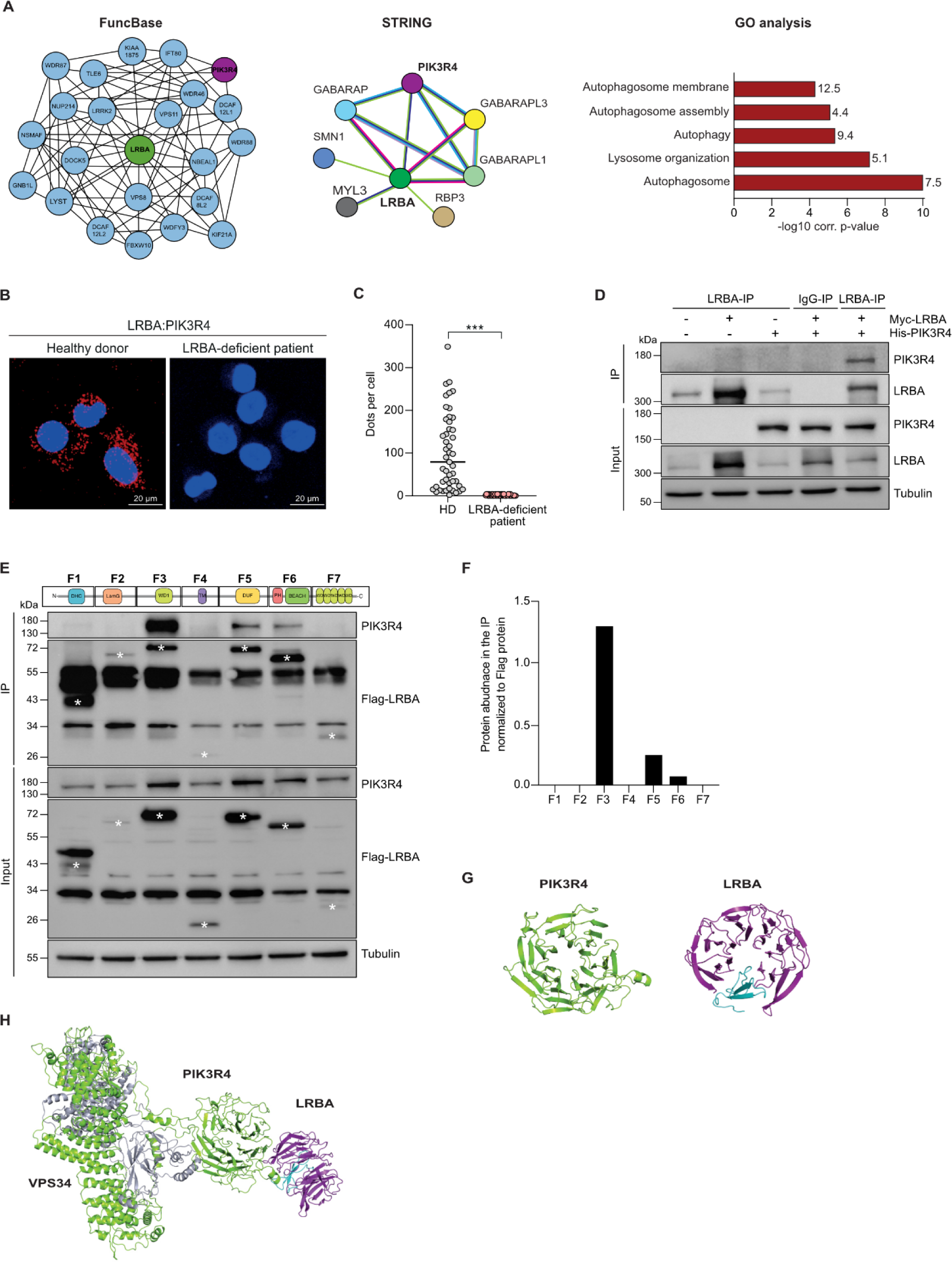
LRBA interacts with PIK3R4 through its WD40 domain. **(A)** *In silico* identification of LRBA interaction protein network using FuncBase (left) and STRING version 10 (middle). In STRING, coloured lines indicate evidence of neighbourhood (green), concurrence (blue), coexpression (black), experimental evidence (purple) and fusion (red). GO enrichment analysis (right) of 28 potential interactors of LRBA was conducted for the three main GO domains cellular compartment. The length of the bar represents the log10 Benjamini-Hochberg corrected p-value. The numbers represent the percentage of associated genes for each biological process. **(B)** Representative fluorescent microscopy images of PLA signal in HD and LRBA-deficient LCL cells. LRBA-PIK3R4 interaction is shown as a red signal and nuclear DAPI staining is shown in blue. Scale bars, 20 µm. **(C)** Dots (PLA signal) per cell were quantified in 50 cells per experiment of HD (grey) and LRBA-deficient (red) LCL cells using Duolink Image tool (SIGMA). A t-test was applied from n=2 independent experiments, ***p<0.001. **(D)** WT HEK293T cells were transfected with a his-tagged PIK3R4 and a myc-tagged LRBA plasmid. Pull down was performed with anti-LRBA and immunoblotted with anti-PIK3R4. **(E)** Schematic representation of the different flag-tagged LRBA plasmids containing the different LRBA protein domains. Each fragment is marked with an asterisk and their corresponding molecular weight are F1: 43 kDa, F2: 65 kDa, F3: 75 kDa, F4: 20 kDa, F5: 68 kDa, F6: 59 kDa, and F7: 31 kDa. HEK293T cells were transfected with his-tagged PIK3R4 and flag-tagged protein fragments 1 to 7 of LRBA. Flag-LRBA pull-downs from transfected cells were immunoblotted with anti-PIK3R4. **(F)** Ratio of PIK3R4/LRBA fragment was calculated for n=2 independent experiments. **(G)** Models of the WD40 domains of PIK3R4 (light green) and LRBA (purple). The inserted repeat propeller of LRBA is shown in cyan. **(H)** Model of the complex of LRBA (purple) with PIK3R4 (light green)/PIK3C3 (VPS34, silver). The inserted repeat propeller of LRBA-WD40 is shown in cyan.

**Table 1.** Predicted LRBA interactors by computational predictions. List of the 28 potential LRBA interactors identified by FuncBase and/or STRING. Proteins are listed alphabetically. Proteins highlighted in grey were selected for validation by co-immunoprecipitations (Co-IP) or proximity ligation assay (PLA) shown in Fig.1 or fig. S1.

| N° | Gen | Protein code | Protein | Function | Database |
| --- | --- | --- | --- | --- | --- |
| 1 | FBXW10 | Q5XX13 | F-box/WD repeat-containing protein 10 | Ubiquitination and subsequent proteasomal degradation of target proteins | FuncBase |
| 2 | DCAF12L1 | Q5VU92 | DDB1-and CUL4-associated factor 12-like protein 1 | Unknown | FuncBase |
| 3 | DCAF12L2 | Q5VW00 | DDB1-and CUL4-associated factor 12-like protein 2 | Unknown | FuncBase |
| 4 | DOCK5 | Q9H7D0 | Dedicator of cytokinesis protein 5 | Epithelial cell migration | FuncBase |
| 5 | GABARAP | O95166 | GABA (A) receptor-associated protein I | Apoptosis and autophagy | STRING |
| 6 | GABARAP L1 | Q9H0R8 | GABA (A) receptor-associated protein like 1 | Formation of autophagosomal vacuoles | STRING |
| 7 | GABARAP L3 | Q9BY60 | GABA (A) receptor-associated protein like 1 | Formation of autophagosomes | STRING |
| 8 | GNB1L | Q9BYB4 | Guanine nucleotide-binding protein subunit beta-like protein 1 | Intracellular signal transduction | FuncBase |
| 9 | IFT80 | Q9P2H3 | Intraflagellar transport protein 80 homolog | Development and maintenance of motile and sensory cilia | FuncBase |
| 10 | KIAA1875 | A6NE52 | WD repeat-containing protein 97 | Generation of catalytic spliceosome | FuncBase |
| 11 | KIF21A | Q7Z4S6 | Kinesin-like protein KIF21A | Microtubule-based movement | FuncBase |
| 12 | LRRK2 | Q5S007 | Leucine-rich repeat<br>serine/threonine-protein kinase | Autophagy through the<br>CaMKK/AMPK | FuncBase |
| 13 | LYST | Q99698 | Lysosomal-trafficking regulator | Sorting endosomal resident<br>proteins into late<br>multivesicular endosomes | FuncBase |
| 14 | MYL3 | P08590 | Myosin light chain 3 | Actin monomer binding | STRING |
| 15 | NBEAL1 | Q6ZS30 | Neurobeachin-like protein 1 | Phospholipid binding | FuncBase |
| 16 | NSMAF | Q92636 | Protein FAN | Ceramide metabolic process | FuncBase |
| 17 | NUP214 | P35658 | Nuclear pore complex protein<br>Nup214 | Nuclear export signal<br>receptor activity | FuncBase |
| 18 | PIK3R4 | Q99570 | Phosphoinositide 3-kinase<br>regulatory subunit 4 | Macroautophagy and late<br>endosome to vacuole<br>transport | FuncBase<br>STRING |
| 19 | RBP3 | P10745 | Retinol- binding protein 3 | Transport of the retinoids | STRING |
| 20 | SMN1 | Q16637 | Survival motor neuron 1 | Assembly of small nuclear<br>ribonucleoproteins | STRING |
| 21 | TLE6 | Q9H808 | Transducin-like enhancer protein 6 | Positive regulation of neuron<br>differentiation | FuncBase |
| 22 | VPS8 | Q8N3P4 | Vacuolar protein sorting-<br>associated protein 8 homolog | Endosomal vesicle fusion | FuncBase |
| 23 | VPS11 | Q9H270 | Vacuolar protein sorting-<br>associated protein 11 homolog | Vesicle-mediated protein<br>trafficking | FuncBase |
| 24 | WDR88 | Q6ZMY6 | WD repeat-containing protein 88 | Unknown | FuncBase |
| 25 | WDFY3 | Q8IZQ1 | WD repeat- and FYVE-containing<br>protein 3 | Aggrephagy | FuncBase |
| 26 | WDR46 | O15213 | WD repeat-containing protein 46 | Scaffold component of the<br>nucleolar structure | FuncBase |
| 27 | WDR87 | Q6ZQQ6 | WD repeat-containing protein 87 | Unknown | FuncBase |

### Proper activity of PI3K-III complex requires LRBA

The PI3K-III complex regulates autophagy by generating PI(3)P, which in turn allows the recruitment of PI(3)P-binding proteins DFCP-1 and WIPI2 to initiate the formation of autophagosomes (*21–24*) (Fig. 2A). Upon autophagy induction with Torin 1 (mTORC1 inhibitor), we observed increased levels of PI(3)P in lipid extracts from WT but not from LRBA-KO HEK293T cells compared to their basal condition (Fig. 2B). Instead, PI(3)P levels in LRBA-KO cells remained similar to those upon inhibition of VPS34 using VPS34-IN1 (Fig. 2B). Moreover, by fluorescence microscopy, we observed significantly reduced accumulation of both WIPI2 and DFCP-1 in LRBA-KO cells in comparison to their WT counterparts after autophagy induction with Torin 1, Rapamycin or upon serum-deprivation (starvation) (Fig. 2, C-E). The specificity of the DFCP-1 signal was validated after VPS34-IN1 treatment (Fig. 2, F and G). Altogether, these results indicate that LRBA is essential for the proper production of PI(3)P and the subsequent recruitment of PI3P-binding effector proteins DFCP-1 and WIPI2 to the forming autophagosome.

**Fig. 2.**
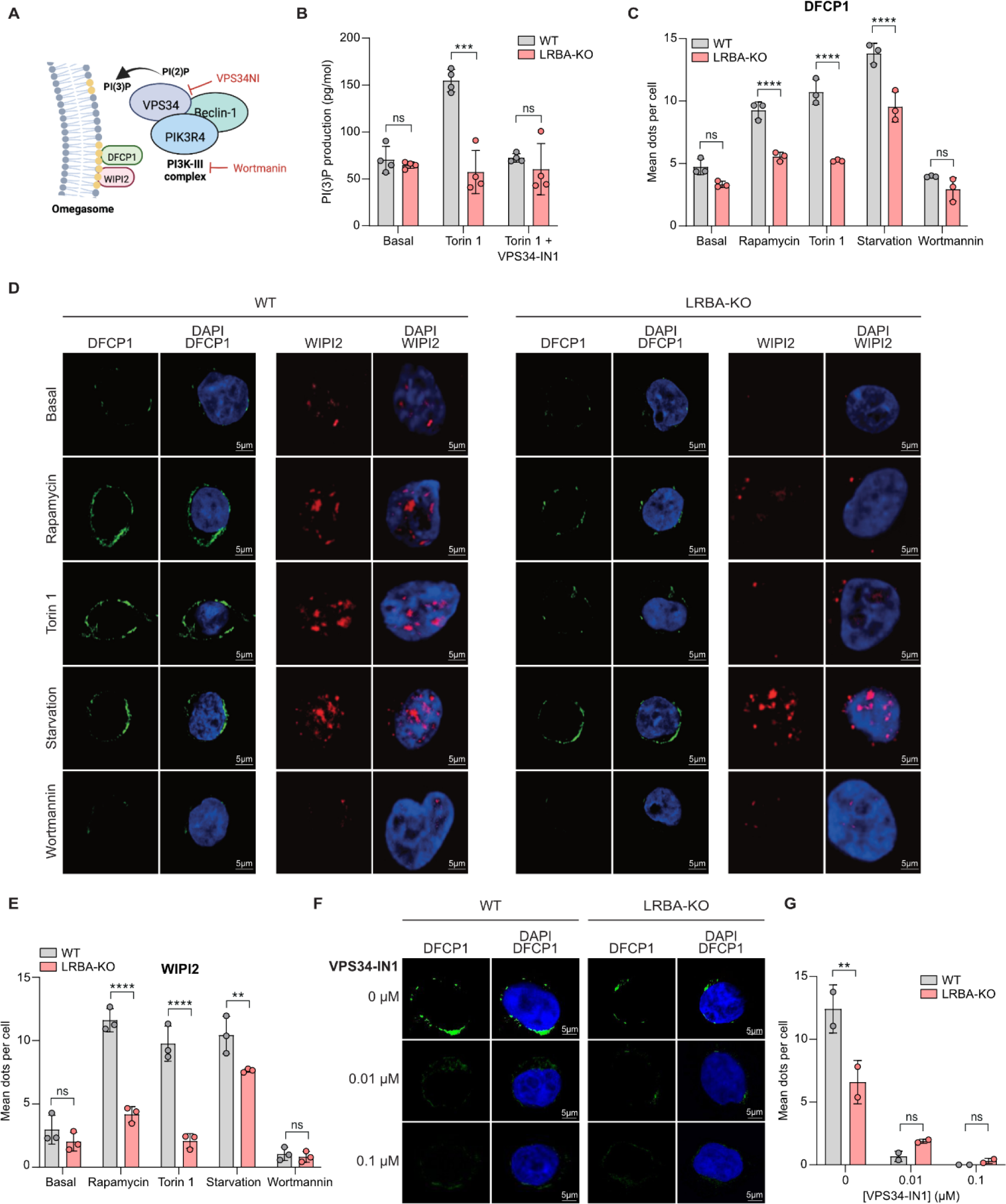
Proper function of the PI3K-III complex requires LRBA. **(A)** Schematic representation of PI3K-III complex activity to initiate autophagosome formation. **(B)** Production of PI(3)P was measured in lipid extracts of WT (grey) and LRBA-KO HEK293T cells (red) under basal conditions, or upon treatment with 333 nM of Torin 1 alone, or in combination with 1 nM of VPS34-IN1 (PI3K-III complex inhibitor) for 4 h. Each dot represents duplicates of one experiment while bars represent the mean ± SD of n=4 independent experiments. A t-test was applied, **p-value <0.01. **(C)** Quantification of DFCP1 signal in WT (grey) and LRBA-KO (red) HEK293T cells stained with anti-DFCP1 antibody after incubation for 1 h with control medium (nuclei: WT=27, KO=21), 50 µM Rapamycin (nuclei: WT=44, KO=49), 333 nM Torin1 (nuclei: WT=52, KO=49), EBSS (nuclei: WT=14, KO=28) or 100 nM Wortmannin (nuclei: WT=29, KO=26). Each dot represents the mean of one experiment while bars represent the mean ± SD of n=3 independent experiments. **(D)** Representative confocal microscopy images of anti-DFCP1 (green) or anti-WIPI2 (red) under different conditions. Scale bar 5 µm. **(E)** Quantification of WIPI2 signal in WT (grey) and LRBA-KO (red) HEK293T cells stained with anti-WIPI2 antibody after incubation for 1 h with control medium (nuclei: WT=59, KO=35), 50 µM Rapamycin (nuclei: WT=70, KO=91), 333 nM Torin1 (nuclei: WT=51, KO=41), EBSS (nuclei: WT=60, KO=42) or 100 nM Wortmannin (nuclei: WT=57, KO=83). Each dot represents the mean of one experiment while bars represent the mean ± SD of n=3 independent experiments. **(F)** Representative confocal microscopy images and **(G)** quantification of WT (grey) (nuclei: 0 μM=46, 0.01 μM=42, 0.1 μM=42) and LRBA-KO (red) (nuclei: 0 μM=29, 0.01 μM=110, 0.1 μM=94). HEK293T cells stained with anti-DFCP1 antibody after treatment with the indicated concentrations of VPS34-IN1 for 1 h followed by 1 h incubation with 333 nM Torin 1. Each dot represents the mean of one experiment while bars represent the mean ± SD of n=2 independent experiments. The statistical analysis of (C), (E) and (G) was performed using a Two-way ANOVA, p-value ****< 0.0001. Scale bar 5 µm.

### LRBA deficiency leads to impaired autophagic flux due to delayed autophagosome-lysosome fusion

We next addressed whether the poor association of WIPI2/DFCP-1 with PI(3)P due to the absence of LRBA affects autophagosome formation, maturation, and ultimately the autophagy flux. Using HEK293T cells stably transduced with an mCherry-GFP-LC3 tandem vector, we were able to differentiate early autophagosomes (green and red signal) and mature autophagolysomes (red signal alone), as the acid-sensitive GFP tag is degraded after lysosomal fusion and LC3 is an autophagosomal marker. We observed higher GFP signal (reduced red/green signal) upon Torin 1 stimulation in LRBA-KO HEK293T cells compared to WT cells (Fig. 3, A and B), suggesting autophagosome accumulation. These observations were confirmed by immunoblotting the conversion of LC3-I into LC3-II in LRBA knockdown (shLRBA) HeLa cells upon stimulation with MG132 and Bafilomycin A1 (fig. S2 A and B) and in LPS-stimulated cells from *Lrba*^-/-^ mice upon stimulation with E64D/Pepstatin A or Bafilomycin A1 (fig. S2, C and D). Interestingly, we observed an increased autophagosomal size based on the LC3 signal evaluation in LRBA-KO HEK293T cells at endogenous levels or after transfection with mCherry-GFP-LC3 (Fig. 3, C-E). Enlarged autophagosomes were also observed in GFP-LC3 transfected shLRBA HeLa cells (fig. S2, E and F). Additionally, we detected accumulation of p62 cargo protein, an autophagy flux marker, in LRBA-KO HEK293T cells (Fig. 3, F and G) and in shLRBA HeLa cells (fig. S2, G and H) upon autophagy induction with Torin1 or MG132, respectively. Taken together our observations indicate a reduced autophagic flux and a deficient cargo degradation due to delayed autophagosome-lysosome fusion in the absence of LRBA.

**Fig. 3.**
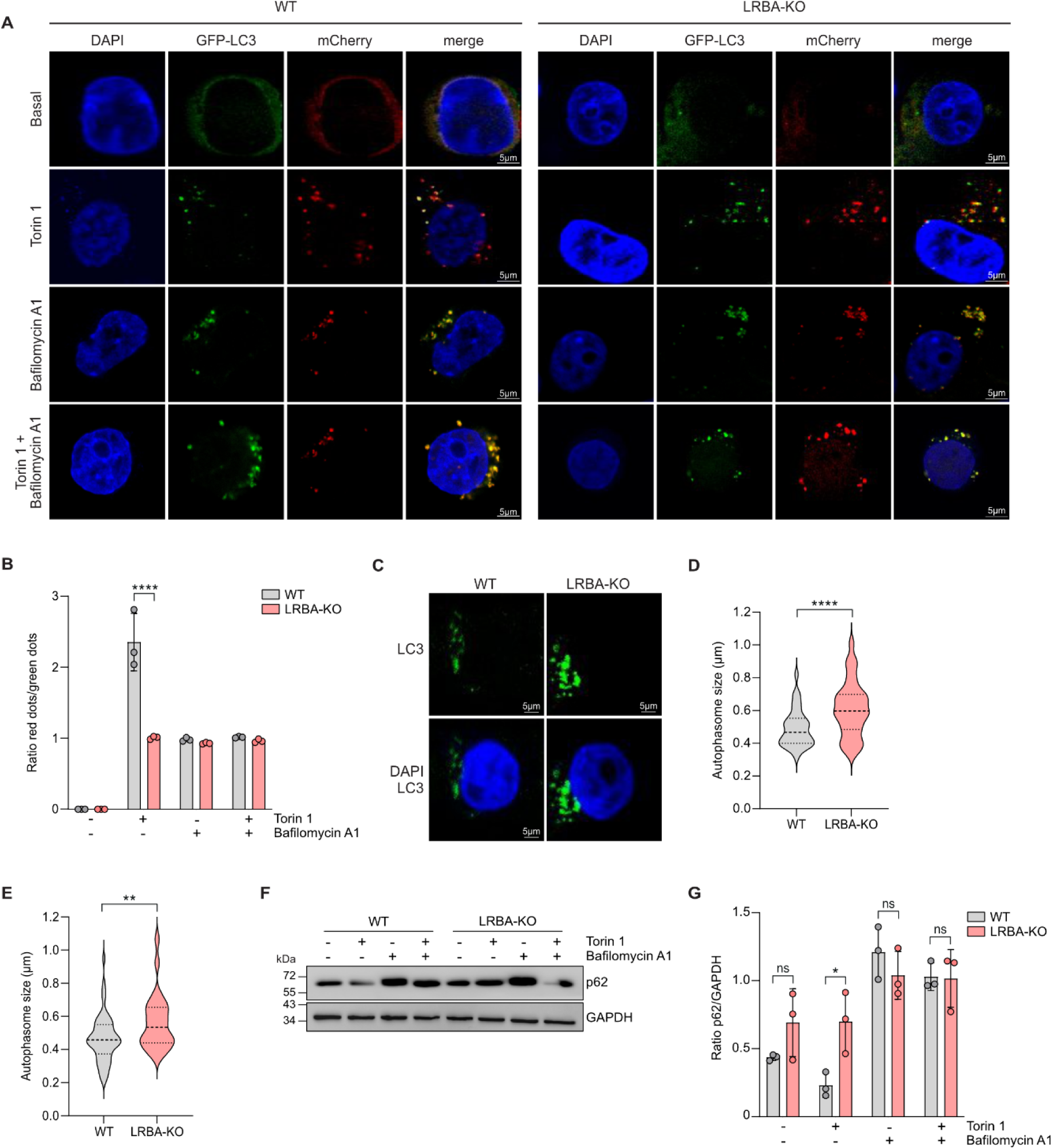
LRBA deficiency leads to impaired autophagic flux due to delayed autophagosome-lysosome fusion. **(A)** Representative confocal microscopy images on WT and LRBA-KO HEK293T cells stably transduced with mCherry-GFP-LC3 tandem vector after 4 h treatment with 333 nM Torin1 alone, 200 nM Bafilomycin A1 alone, or in a combination of both. **(B)** Ratio red/green dots counted with ImageJ in WT (grey) (nuclei: Torin1=87, Bafilomycin A1=86, Torin1+Bafilomycin A1=86) and LRBA-KO HEK293T (red) (nuclei: Torin1=81, Bafilomycin A1=93, Torin1+Bafilomycin A1=84). Each dot represents the mean of one experiment while bars represent the mean ± SD of n=3 independent experiments. Statistical analysis was performed using a Two-way ANOVA, p-value ****< 0.0001. **(C)** Representative confocal microscopy images of LC3 signal at endogenous levels upon Bafilomycin A1 stimulation for 1 h using anti-LC3 antibody in WT and LRBA-KO HEK293T cells. **(D)** Measurement of autophagosome diameter (in µm) of (C) (n=68 autophagosomes) in WT (grey) and LRBA-KO (red) HEK293T cells. The dashed lines represent the mean and the dot lines represent the first and third quartiles. **(E)** Measurement of autophagosome diameter (µm) in mCherry-GFP-LC3 transduced WT (grey) and LRBA-KO (red) HEK293T cells (n=50 autophagosomes). **(F)** Representative immunoblot analysis of p62 expression in WT and LRBA-KO HEK293T cells at basal conditions or after 4 h stimulation with 333 nM Torin 1 alone, with 200 nM Bafilomycin A1 alone, or with a combination of both. GAPDH was used as housekeeping protein. **(G)** Densitometry analysis of p62 immunoblots in WT (grey) and LRBA-KO HEK293T cells (red). Bars represent the mean ± SD from n=3 independent blots. Statistical analysis was performed using Welch’s test p=0.0017 (*).

### Loss of LRBA leads to abnormal accumulation of enlarged, sealed and unfused autophagosomes

To analyse whether these enlarged autophagosomes are matured and sealed, we performed a protease protection assay and monitored the sensitivity of p62 cargo protein to proteinase K treatment (Fig. 4A). We observed increased p62 protease protection in LRBA-KO cells compared to WT HEK293T cells, suggesting an accumulation of sealed autophagosomes in the absence of LRBA (Fig. 4, B and C). Accumulation of enlarged autophagosomes was confirmed by membrane flotation on OptiPrep gradients (*25*). With this method, unfused autophagosomes float at 5% OptiPrep, whereas autophagic precursor membranes, ER and lysosomes float at higher densities. In contrast to WT cells, we found increased LC3-II signals in 5% fractions in LRBA-KO cells under starvation, suggesting an accumulation of unfused autophagosomes. Following autophagosome enrichment using Bafilomycin A1, LC3-II appeared in lower densities of 5%, indicating the formation of mature autophagosomes (Fig. 4D). Similar results were obtained when monitoring the autophagosomal soluble N-ethylmale-imide-sensitive factor-attachment protein receptors (SNARE) protein STX17 (Fig. 4D). Accumulation of enlarged autophagosomes was also observed by EM after LC3 immunogold labelling in isolated LPS-stimulated B cells from *Lrba*^-/-^ mice (Fig. 4E). In WT mice, however, autophagosomes were hardly detectable due to their rapid turnover. Taken together, loss of LRBA affects the biology of autophagosomes, resulting in the accumulation of enlarged, sealed autophagosomes that fuse less with lysosomes.

**Fig. 4.**
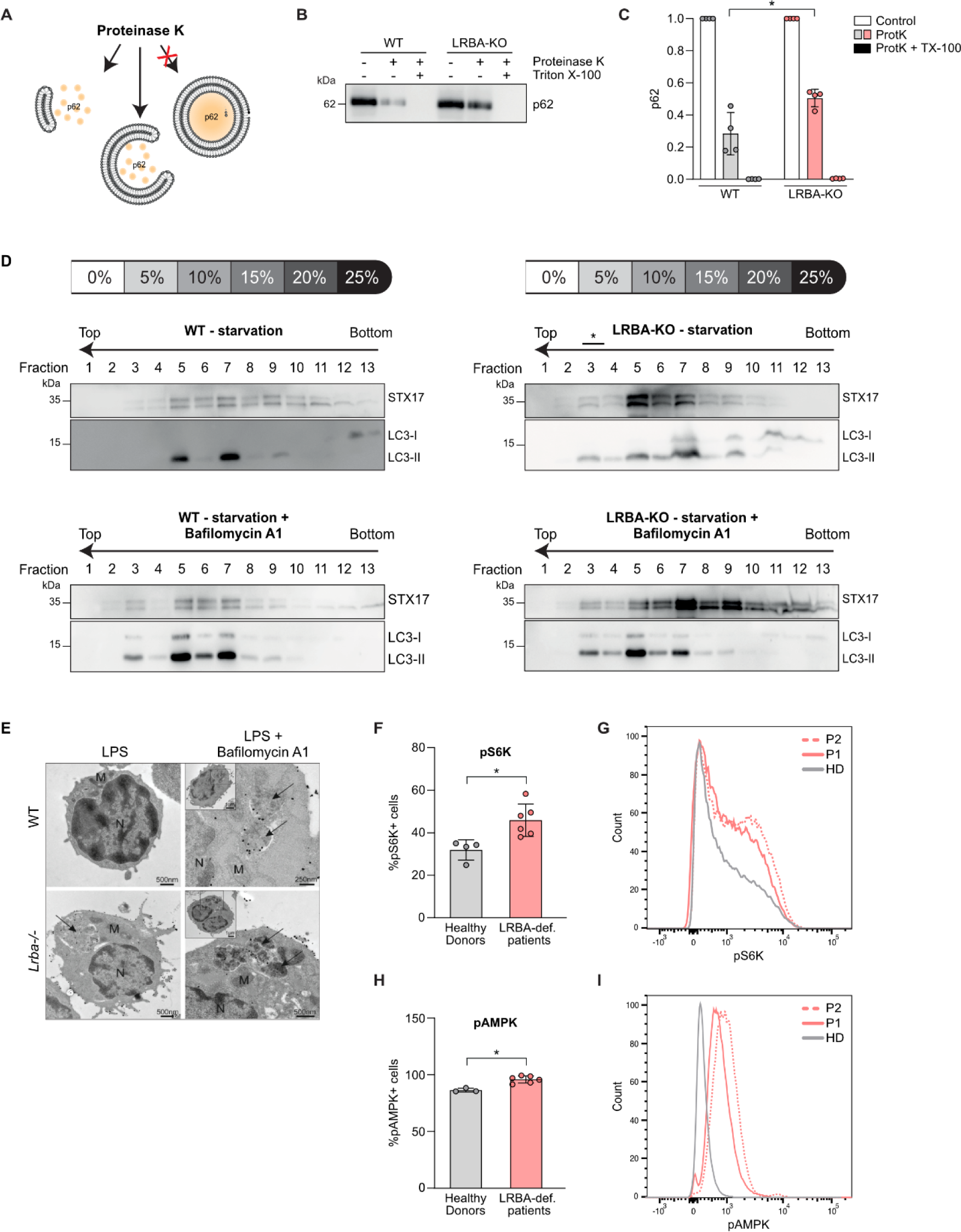
Loss of LRBA leads to abnormal accumulation of enlarged, sealed and unfused autophagosomes and to increased mTOR signaling. **(A)** Schematic illustration of the protease protection assay. Cell homogenates are treated with proteinase K resulting in the degradation of exposed proteins including p62 but not of membrane-enclosed proteins. **(B)** WT and LRBA-KO HEK293T cells were grown in EBSS medium with 300 nM Torin 1 for 2 h. After lysis, cell membranes were subjected to proteinase K and Triton X-100 treatment as indicated and analyzed by anti-p62 Western blotting. **(C)** Quantification of (B) in WT (grey) and LRBA-KO (red) HEK293T cells. Bars represent the mean ± SEM from n=4 independent blots. Statistical analysis was performed using Welch’s t-test p-value *<0.05. **(D)** WT and LRBA-KO HEK293T cells were grown in EBSS medium with 300 nM Torin1 for 3 h with or without 200 nM Bafilomycin A1, and cell homogenates were subjected to OptiPrep flotation analysis. The asterisk indicates the accumulation of autophagosomes in LRBA-KO homogenates. **(E)** Electron microscopy images on LPS-stimulated B cells from WT and *Lrba^-/-^* mice using immunogold labeling for LC3. Arrows point to autophagosomes, N: nucleus, M: mitochondria. Images represent higher magnification of the inset’s black-boxed area. A total of three mice per genotype were analyzed. Scale bars=500 nm for LPS WT and *Lrba-/-* and LPS+Bafilomycin A1 for *Lrba-/-*, 250 nm for LPS+Bafilomycin A1 WT. **(F)-(I)** Flow cytometry analysis showing increased expression of pS6K and pAMPK in LCL from LRBA-deficient patients. **(F), (H)** Bar graphs represent the mean percentage of LCL from HD (grey) and LRBA-deficient patient (red) positive for pS6K **(F)** or for pAMPK **(H)** from n=3 independent experiments. Representative histogram of the MFI of **(G)** pS6K or **(I)** in HD (grey) and two LRBA-deficient patients (red). Welch’s test was applied, p=0.0073 (**) and p=0.0007 (**).

### Increased mTORC1 signalling in the absence of LRBA

The PI3K-III complex regulates the mechanistic/mammalian target of rapamycin complex (mTORC1) (*26, 27*), which in turn regulates autophagy. We therefore evaluated the phosphorylation status of the mTOR *bona fide* substrate, p70-S6K1 (*28*). Hyperphosphorylation of p70-S6K1 was observed in LCL cells from two LRBA-deficient patients and in shLRBA HeLa cells (Fig. 4, F and G; fig. S2, I and J). AMP-dependent kinase (AMPK) also showed increased activity, as assessed by phosphorylation at T172 in both cell lines (Fig. 4, H and I; fig. S2, I and J). AMPK inactivates mTORC1 and enhances autophagy (*29*) and may thus, act to sustain residual autophagy, compensating for the decreased activity of the PI3K-III complex in the absence of LRBA. Together, these observations suggest a role of LRBA in the regulation of mTORC1 activation.

### LRBA interacts with FYCO1 facilitating autophagosome mobilization and lysosome positioning

To explain the decreased cargo elimination *via* autophagy, we evaluated autophagosome mobility by live-microscopy using GFP-LC3 transfected HEK293T cells. We observed that the enlarged autophagosomes in LRBA-KO cells have reduced velocity in comparison to regular-sized autophagosomes (Fig. 5, A-C and supplementary video 1 and 2). Interestingly, lysosomal positioning was altered in LRBA-KO cells, showing a cytosolic spread-out signal in contrast to the perinuclear location in WT cells (Fig. 5D). However, LRBA-KO cells showed normal lysosome biogenesis evidenced by endogenous Transcription Factor EB (TFEB) signals (Fig. 5E). In addition, EM analyses showed lysosome accumulation in B-lymphocytes from *Lrba*^-/-^ mice upon LPS *in vitro* stimulation (Fig. 5F). Together, these data suggest a role of LRBA in autophagosome and/or lysosome trafficking. To confirm these observations, we searched for another LRBA interaction partner specifically involved in autophagosome and lysosome trafficking using unbiased IP-MS/MS data previously reported (*30*). In this work, LRBA is found in the FYCO1 interactome (http://besra.hms.harvard.edu/ipmsmsdbs/comppass_viewer.php?bait=FYCO1). FYCO1 is a multidomain adaptor protein that mediates microtubule plus-end-directed autophagosome transport by simultaneously interacting with kinesin motor proteins and autophagosomal proteins such as Rab7, LC3 and PI(3)P (*14, 15, 31*). Following co-IP analysis in HEK293T cells co-transfected with GFP-tagged FYCO1 and myc-tagged LRBA, we confirmed a physical interaction between LRBA and FYCO1 (Fig. 5G). Additional co-IP experiments using seven different flag-tagged LRBA protein domains (F1-F7) and GFP-tagged FYCO1, identified several potential binding sites contained in domains F1, F5 and F6 (Fig. 5, H and I). Altogether, our results show that LRBA interacts with PIK3R4 and FYCO1 under basal autophagy conditions, suggesting a role of LRBA at different autophagy stages by forming different protein complexes.

**Fig. 5.**
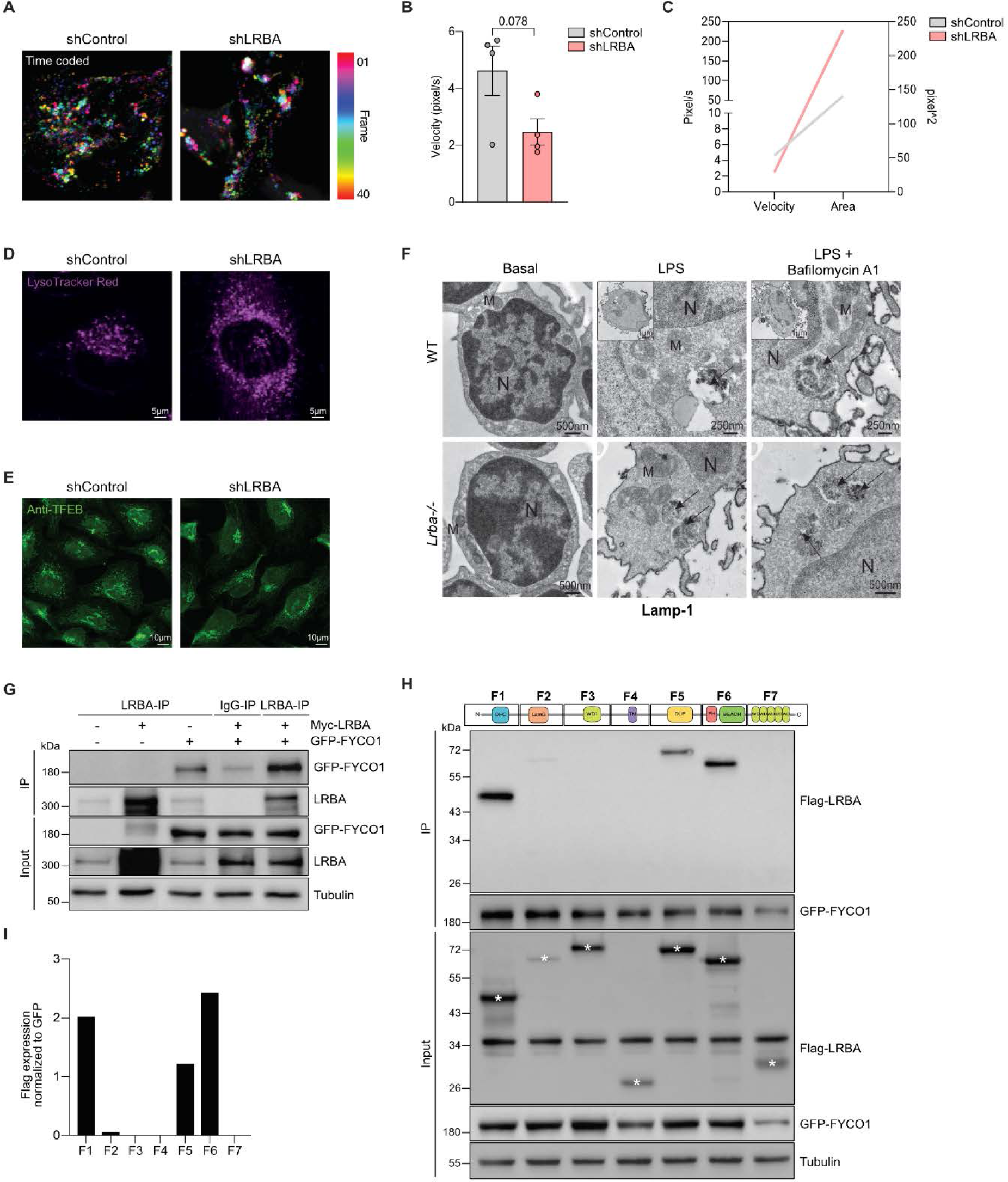
LRBA-FYCO interaction facilitates autophagosome mobilization and lysosome positioning. **(A)** Mobility of autophagosomes was evaluated by time-lapse movies (supplementary video 1 and 2) acquired from individual frames, color-coded and projected into one image, using the temporal color-coding module of the Zeiss Zen Black software. **(B)** Quantification of autophagosomes velocity was performed using Manual Track. Each dot corresponds to one autophagosome tracked over 25 frames for shControl (grey) and shLRBA HeLa cells (red). Bars represent mean ± SD from n=4 tracked autophagosome. Welch’s t-test was applied. **(C)** Correlation graph of velocity and area of the autophagosomes tracked in shControl (grey) and shLRBA HeLa cells (red). **(D)** Lysosomes from shControl and shLRBA HeLa cells under basal conditions were stained with LysoTracker red. shLRBA HeLa cells showed cytosolic spread-out of lysosomes in comparison to perinuclear lysosomal distribution in the shControl cells. **(E)** Representative fluorescent microscopy images of shControl and shLRBA cells stained with anti-TFEB antibody upon resting conditions showing normal biogenesis of lysosomes in absence of LRBA**. (F)** Electron microscopy images on LPS-stimulated B cells from WT and *Lrba^-/-^* mice using immunoperoxidase labeling for Lamp-1. Arrows point to lysosomes. N: nucleus, M: mitochondria. Images represent higher magnification of the inset’s black-boxed area. A total of three mice per genotype were analyzed. Scale bars=500 nm for WT Basal and *Lrba-/-* Basal, LPS, LPS+Bafilomycin A1 and 250 nm for WT LPS, LPS+Bafilomycin A1. **(G)** WT HEK293T cells were transfected with DNA encoding GFP-FYCO1 and/or Myc-LRBA plasmid. Pull down was performed with anti-LRBA and immunoblotted with anti-GFP. **(H)** WT HEK293T cells were transfected with DNA encoding GFP-FYCO1 plasmid and flag-tagged fragments 1 to 7 of LRBA. Flag-LRBA immunoprecipitates from transfected cells were immunoblotted with anti-GFP. **(I)** Ratio of Flag expression normalized to GFP expression of n=2 independent blots.

### Enhanced T-cell response due to increased MHC class II restricted antigen presentation in LRBA deficiency

Autophagy delivers self and foreign cytoplasmic proteins to lysosomes for further loading onto MHC-II molecules. MHC-II-peptide complexes are transported to the cell membrane for antigen presentation to CD4^+^ T lymphocytes (*32*). To test whether defective autophagy due to LRBA loss affects MHC-II mediated antigen presentation, we used the Influenza A matrix protein 1 (MP1) fused to LC3 (MP1-LC3), which enhances its presentation *via* MHC-II to CD4^+^ T-cell clones (*11*). Thus, the abilities of WT and LRBA-KO HaCat cells expressing MP1-LC3 (target cell) to activate MP1-specific CD4^+^ T-cells (effector cells) were assessed by their cytokine secretion in response to this stimulation (Fig. 6A). We detected an increase of IFN-γ and IL-6 secretion in the supernatant of CD4^+^ T-cells co-cultured with LRBA-KO HaCat cells compared to WT HaCat cells (Fig. 6, B and C). Secretion of TNF-α and MIP-1β were also found increased in the supernatant of LRBA-KO HaCat cells (fig. S3, A and B). This suggests increased MHC-II antigen presentation through autophagy, enhancing CD4^+^ T-cell stimulation in the absence of LRBA. Further confocal microscopy analysis of the MHC-II compartment (MIIC) for antigen loading revealed an enlarged size and increased number of HLA-DR or HLA-DR/LC3 containing vesicles in LRBA-KO HaCat cells (Fig. 6, D-F). In addition, we observed a trend towards increased LC3/HLA-DR co-localization in LRBA-KO HaCat cells (Fig. 6G). This suggests an accumulation of autophagosomes delivering antigens inside the MIICs of LRBA-deficient cells. These observations support our findings of enlarged autophagosomes and accumulation of p62 cargo protein in LRBA-KO HEK293T cells. Altogether, our data suggest that lack of LRBA results in increased MHC-II restricted antigen presentation of autophagy substrates by delaying the maturation and fusion of autophagosomes with lysosomes.

**Fig. 6:**
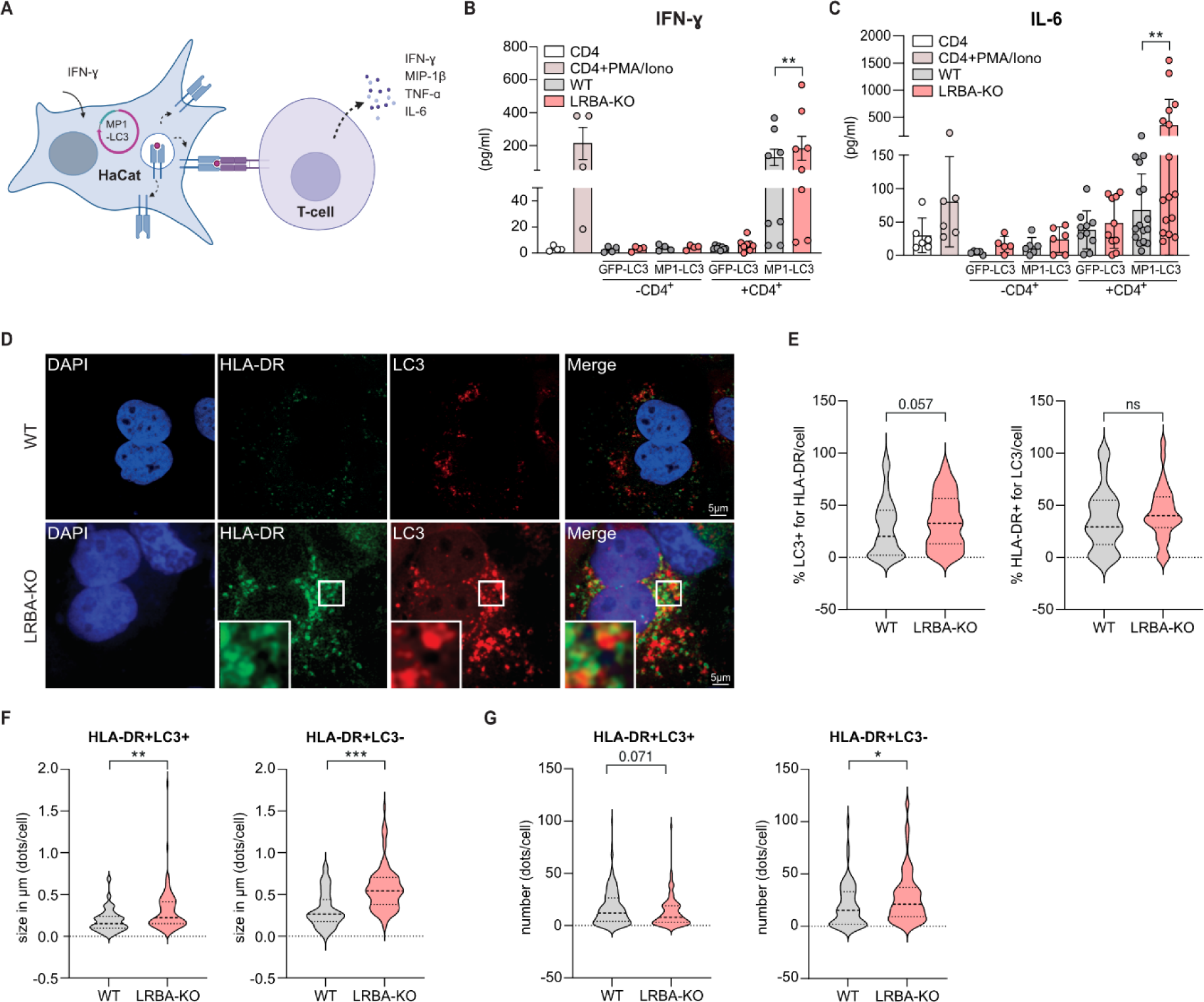
Enhanced T-cell response due to increased MHC class II restricted antigen presentation in LRBA deficiency. **(A)** Schematic illustration of the antigen presentation assay. WT or LRBA-KO HaCat cells stably expressing LC3-GFP or MP1-LC3 (target cells), and pre-treated with IFN-γ for 24 h to up-regulate MHC-II molecules were cultured with a MP1-specific CD4^+^ T cell clone (effector cells) from a HD for 20 h. Following incubation, proinflammatory cytokines were measured in the culture supernatants. **(B)-(C)** ELISA assays on culture supernatants from co-cultures of an T-cell effector to HaCat target cell (E:T=10:5) ratios as described in A. Bars represent the mean ± SD of **(B)** IFN-γ (n=4) and **(C)** IL-6 (n=4) secretion in WT (grey) and LRBA-KO (red) HaCat cells and each dot represents a replicate. Ratio paired student t-test, Two-tailed: p-value *<0.05, **<0.0005. **(D)** Representative confocal microscopy images of HaCat cells stimulated overnight with IFN-γ and treated with 20 µM chloroquine for 6 h. Fixed cells were immunostained for HLA-DR (green), LC3 (red) and DAPI (blue) for the nuclei and then acquired using a SP8 Falcon confocal microscope (Leica), scale bar 5 µm. **(E)** Graph represents the percentage of LC3^+^ dots also positive for HLA-DR per cell (left) and the right panel shows the percentage of HLA-DR dots also positive for LC3 per cell in WT (grey) and LRBA-KO (red) HaCat cells with more than 30 cells quantified per condition. Quantification of LC3, HLA-DR or double positive dots per cell was performed using a semiautomatic plugin designed on FIJI. The violin blot shows the average size in µm **(F)** or the number **(G)** of dots positive for HLA-DR per cell in WT (grey) and LRBA-KO (red) HaCat cells with more than 30 cells quantified per condition. Statistical analysis was performed using unpaired t-test. p-values *< 0.05, **<0.01, ***< 0.0001.

### LRBA reconstitution restores DFCP-1 recruitment, p62 degradation, and number and size of HLA-DR/LC3 vesicles

To further link the lack of LRBA to the observed abnormal autophagy, LRBA-KO HEK293T cells and LRBA-KO HaCat cells were reconstituted for LRBA expression by transitional transfection with a myc-tagged WT-LRBA plasmid (hereafter called Myc-LRBA) or an empty vector (mock control) (fig. S4, A and B). Reconstituted Myc-LRBA cells were tested for autophagosome formation (DFCP-1 recruitment), autophagy flux (p62 degradation) and antigen presentation (HLA-DR/LC3 vesicles), which were further compared to WT and LRBA-KO cell lines. Following autophagy induction by Torin 1 and using fluorescence microscopy, we observed a 3.9-fold increase of DFCP-1 expression in Myc-LRBA reconstituted HEK293T cells compared to LRBA-KO cells (Fig. 7, A and B). In addition, the autophagy flux marker, p62, was assessed by immunoblotting under basal conditions or upon Torin 1 stimulation. We observed a significant decrease of p62 accumulation in Myc-LRBA reconstituted cells in comparison to LRBA-KO cells (Fig. 7, C and D). Together, our data indicate that ectopic expression of LRBA restores the recruitment of DFCP-1 and the degradation of p62 suggesting a normalization of autophagosome formation and autophagy flux observed in absence of LRBA. In addition, we observed a significant reduction of HLA-DR/LC3 vesicle number and size upon IFN-γ stimulation in Myc-LRBA reconstituted HaCat cells in comparison to LRBA-KO cells (Fig. 7, E-G). Similar number and size of HLA-DR/LC3 vesicles were observed in the Myc-LRBA and WT conditions, suggesting a reduction of autophagy substrates accumulation in MIICs (Fig. 7, E-G). Reduced expression of HLA-DR molecules upon IFN-γ stimulation in Myc-LRBA HaCat cells was further confirmed by flow cytometry (fig. S4, C and D). Finally, HLA-DR/LC3 colocalization was also rescued to WT levels after ectopic expression of LRBA (Fig. 7H). Together these results directly implicate LRBA in the autophagy-dependent antigen presentation.

**Fig. 7.**
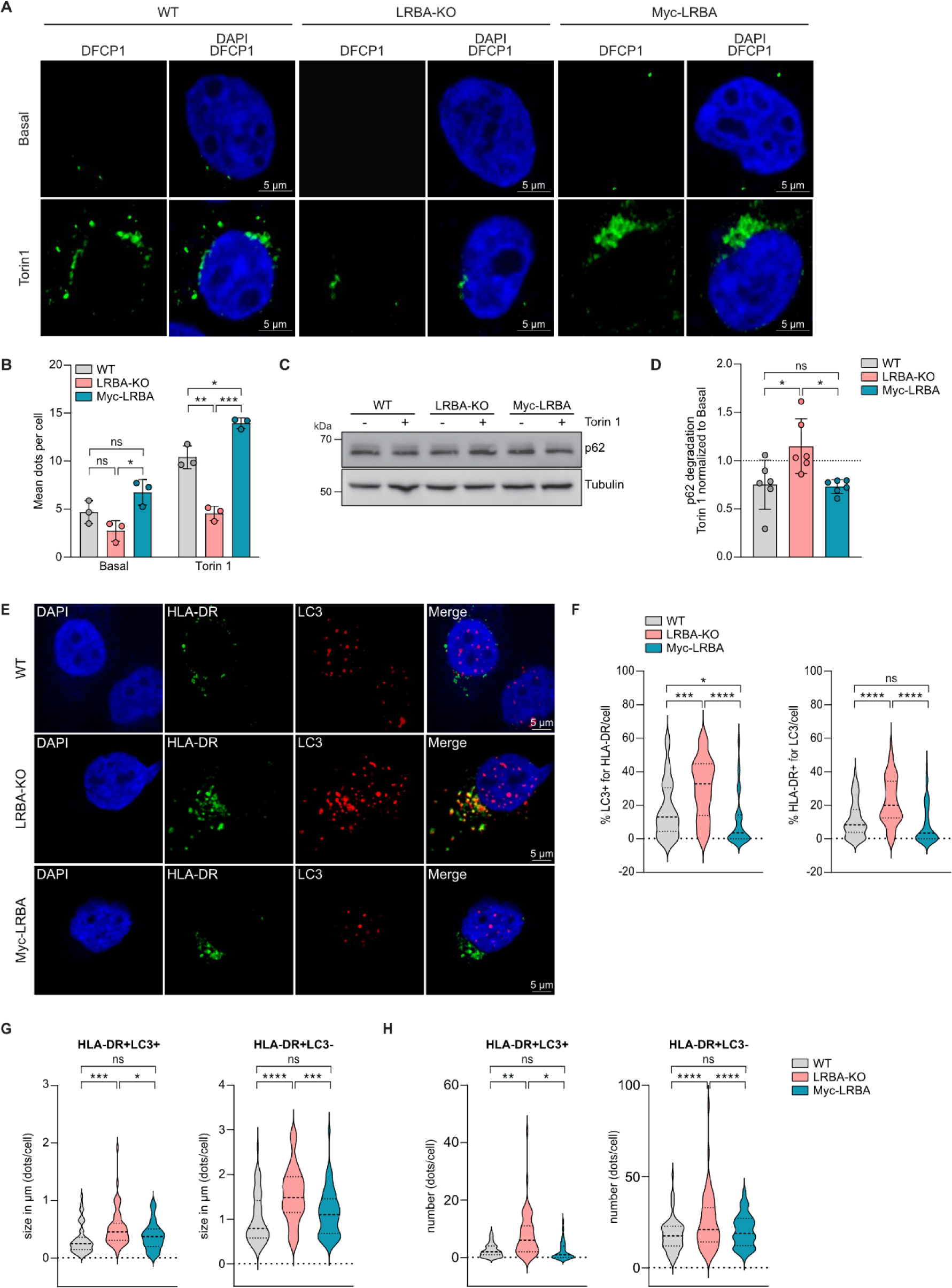
LRBA reconstitution restores DFCP-1 recruitment, p62 degradation and number and size of HLA-DR/LC3 vesicles. **(A)-(B)** DFCP-1 expression in WT, LRBA-KO and Myc-LRBA HEK293T cells after stimulation for 1 h with control medium or 333 nM Torin1. **(A)** Representative confocal microscopy images. Scale bar 5 µm. **(B)** Quantification of DFCP-1 signal by confocal microscopy using anti-DFCP1 in WT (grey) (nuclei: basal=158, Torin 1= 244), LRBA-KO (red) (nuclei: basal=269, Torin 1=294) and Myc-LRBA (teal) HEK293T cells (nuclei: basal=152, Torin 1=273.Each dot represents the mean of one experiment while bars represent the mean ± SD of n=3 independent experiments. **(C)** Representative immunoblot analysis of p62 and Tubulin expression in WT, LRBA-KO and Myc-LRBA HEK293T cells at basal conditions or after 4 h stimulation with 333 nM Torin 1. **(D)** Densitometry analysis of p62 degradation after Torin 1 stimulation normalized to basal conditions of WT (grey), LRBA-KO (red) and Myc-LRBA (teal). Bars represent the mean ± SD of n=6 independent blots. **(E)** Representative confocal microscopy images of WT, LRBA-KO and Myc-LRBA HaCat cells stimulated overnight with IFN-γ and treated with 20 µM chloroquine for 6 h. Fixed cells were immunostained for HLA-DR (green), LC3 (red) and DAPI (blue) and then acquired using a LSM 710 (Zeiss). Scale bar 5 µm. **(F)** Graph represents the percentage of LC3^+^ dots that are also positive for HLA-DR per cell (left; 60 nuclei) and the percentage of HLA-DR^+^ dots that are also positive for LC3 per cell (right; 60 nuclei) in WT (grey), LRBA-KO (red) and Myc-LRBA (teal) HaCat cells. **(G)-(H)** Violin plots represent the size **(G)** and number **(H)** of dots positive for LC3 and also positive for HLA-DR per cell (left; 60 nuclei) and the dots positive for HLA-DR dots also positive for LC3 per cell (right; 60 nuclei) in WT (grey), LRBA-KO (red) and Myc-LRBA (teal) HaCat cells. Quantification of LC3, HLA-DR or double positive dots per cell was performed using a semiautomatic plugin designed on FIJI. n=3 independent experiments. Welch’s t-test was applied for (B), (D), (F)-(H), p-values *< 0.05, **<0.01, ***< 0.0001, **** < 0.0001.

## Discussion

In this study, we propose a novel role of LRBA as a scaffold protein required at early and late stages of autophagy through the binding to PIK3R4 and FYCO1. Through these interactions, LRBA facilitates the proper production of PI(3)P upon autophagy induction, as well as the formation of autolysosomes required for the degradation of the sequestered cargo material. Through this mechanism, LRBA regulates MHC-II restricted antigen presentation and T-cell mediated responses.

While our proposed mechanism stands as a crucial factor in understanding the hyperinflammation seen in LRBA deficiency, it is not the sole responsible pathomechanism. Instead, it serves as an additional independent mechanism, distinct from e.g. CTLA4 trafficking, that worsens the inflammatory response in T cells, contributing to an increased severity, elevated frequency of enteropathy, higher disease mortality, and earlier onset of LRBA deficiency when compared to CTLA4 insufficiency (*7, 33*). In fact, human inflammatory bowel disease (IBD) pathogenesis, one of the most prominent clinical manifestations in LRBA deficiency, has been clearly linked to abnormal autophagy (*34, 35*). However, future analysis addressing additional implications of disrupted autophagy in other autophagy-dependent biological mechanisms, or addressing the functionality of other types of autophagy (such as mitophagy, or ER-phagy) in other cell models, will contribute to a comprehensive understanding of the multi-faceted roles of LRBA. In fact, a recent study has shown that LRBA regulates ATG9A vesicle trafficking in HeLa cells and absence of LRBA resulted in reduced mitophagy efficiency (*36*).

We found that LRBA-PIK3R4 interaction occurs *via* WD40-WD40 domain binding (*37*). WD40 domains are characteristically present in BDCP family members like LRBA, allowing them to act as scaffolding proteins, or like WDFY3/ALFY enabling its interaction with PI3K-III complexes to participate in cargo selection and delivery to autophagosomes (*38, 39*). In contrast, WD40 domains are not abundant in PI3K family members, only PIK3R4 harbours a WD40 domain within this protein family, supporting that LRBA specifically interacts with this protein and not with other PI3K family members (*40*). In addition, LRBA interacts with FYCO1 *via* its BEACH domain, a domain involved in vesicle trafficking of CTLA-4-containing vesicles in Tregs (*6*). This indicates that LRBA forms distinct protein complexes through its BEACH and WD40 domains, playing a role in both CTLA-4 mobilization (*6*) and autophagy. LRBA might also act in additional vesicle trafficking pathways forming different protein complexes. Future analyses could address whether LRBA protein interactions are cell specific or stimuli specific.

We observed that the LRBA-PIK3R4 interaction facilitates the production of PI(3)P, as LRBA-KO cells failed to produce PI(3)P after autophagy induction. At resting conditions, however, both WT and LRBA-KO cells produced equal amounts of PI(3)P suggesting that the PI(3)P constitutive pool is not affected by the loss of the LRBA-PIK3R4 interaction. Indeed, downregulation of VPS34 expression in resting glioblastoma cells resulted in a residual production of PI(3)P (*41*), indicating that the PI(3)P constitutive pool is generated independently of the PI3K-III complex. (*41–43*). VPS34-KO cells showed normal vesicle trafficking between the *trans*-Golgi and late endosomes, normal endocytic uptake of fluid-phase markers, and normal association with early endosomes, but disrupted late endosomal trafficking (*27*). Therefore, the effect of LRBA in CTLA-4 vesicle trafficking could also be related to an aberrant activity of the PI3K-III complex. Moreover, we demonstrated that LRBA interacts with UVRAG, which together with PIK3R4 and VPS34 forms a protein complex located at the membrane of early endosomes acting in endocytosis, cytokinesis and lysosome recycling, suggesting that LRBA might also affect other vesicle trafficking processes.

PI(3)P acts as a membrane-bound localized signal, controlling the assembly of PI(3)P-binding scaffold proteins such as WIPI2 and DFCP-1 that mediate autophagosome biogenesis (*44*). WIPI2 is necessary for recruiting the lipidation machinery for the phagophore-forming protein LC3B (*45, 46*), whereas the ATPase DFCP1 is necessary for releasing the newly formed autophagosomes from the ER (*47*). Despite the reduction of WIPI2 and DFCP1 recruitment upon autophagy induction in LRBA-KO cells, comparable levels of LC3 lipidation to WT cells were observed. Normal autophagosome formation was also seen in DFCP1-mutant cells and in Vps34-mutant cells, despite reduced WIPI2 puncta (*48*). Similar to our observations, shVPS34 glioblastoma cells as well as *vps34*-KO MEF cells showed enlarged late endosomes but with an intact capacity to fuse with lysosomes (*41*). Our data, however, shows a delay of autophagosome/lysosome fusion in LRBA-KO cells, which highlights the additional role played by LRBA in the autophagy process. We suggest that the LRBA-FYCO1 interaction is essential for autophagosome mobilization and regular lysosome positioning followed by autophagosome/lysosome fusion. This notion is reinforced by previous reports showing the importance of FYCO1 for intracellular transport of autophagic vesicles (*14, 15, 49*), lysosomes and phagosomes (*50, 51*). In fact, FYCO1 depletion led to accumulation of autophagosomes as observed in LRBA-KO cells (*30*). Since ectopic expression of Myc-LRBA in LRBA-KO HEK293T cells rescued the expression of DFCP1 and the degradation of p62, our data support the requirement of LRBA for autophagosome maturation and therefore autophagy flux.

Autophagy is essential for multiple immune functions including cell survival and differentiation, pathogen clearance, cytoskeleton polarization, and antigen processing for MHC-II restricted presentation. We have previously reported defective autophagy in naïve B cells from LRBA-deficient patients, which was associated with poor cell survival, reduced plasmablast differentiation and low antibodies production (*1*). The importance of autophagy for plasmablast survival and differentiation has been demonstrated in Atg5-deficient mice (*52, 53*). In T lymphocytes, the PI3K-I complex has been revealed to be the mayor autophagy player upon starvation and TCR stimulation, whereas PI3K-III complex activity has been shown to be completely dispensable (*54*). In contrast, it has also been shown that PI3K-III complex dependent autophagy is required for naïve T-cell homeostasis through the quality control of mitochondria (*55*). Together these observations may explain the impaired activation of mTORC1 and mTORC2 complexes in human LRBA-deficient Tregs (*via* PI3K-I complex) and the skewed T-cell memory phenotype (*via* PI3K-III complex) in LRBA deficiency (*56*). Although abnormal autophagy seems not to affect T-cell survival, as LRBA-deficient patients have normal numbers of circulating T-cells, it might affect endocytic trafficking routes. Previous findings showed an essential role of autophagy in MHC-II restricted presentation to CD4^+^ T-cells using HaCat cells transfected with MP1-LC3 plasmid (*11*). Using this setup in LRBA deficiency, HaCat cells showed enlarged vesicles with increased co-localization of LC3 and the MHC-II molecules HLA-DR, suggesting an accumulation of autophagy substrates in MIICs. Similar vesicle enlargement and accumulation of the autophagy cargo protein p62 was also observed in LRBA-KO HEK293T and shLRBA HeLa cells, confirming that the absence of LRBA leads to an accumulation of autophagosomes. Rescue experiments based on ectopic expression of WT-LRBA showed decreased co-localization of HLA-DR/LC3 vesicles with levels similar to those observed in WT cells. Furthermore, MP1-specific CD4^+^ T-cell clones co-cultured with cells lacking LRBA resulted in increased proinflammatory cytokine secretion, indicating enhanced activation of CD4^+^ T-cells in the absence of LRBA. Accumulation of enlarged MIICs suggests that the missing interaction of LRBA and FYCO1 can block the trafficking of HLA-DR containing vesicles, allowing prolonged peptide loading onto MHC-II. With time the autophagy delivered peptide repertoire will be processed, leading to a prolonged antigen presentation that results in enhanced and chronic T-cell activation. These observations are in line with the clinical picture of LRBA-deficient patients, which is characterized by T-cell immune dysregulation (*2, 4, 5*). An enhanced proinflammatory response in the context of LRBA deficiency has also been reported for *Lrba^-/-^* mice that display increased susceptibility to DSS-induced colitis. This has been attributed to abundant production of type I IFN in IRF3/IRF7-and PIK/mTOR-dependent signalling in dendritic cells, thus recognizing a role of LRBA in the endolysosomal pathway to limit TLR signalling (*57*).

Finally, we show that LRBA-deficient LCL cells exhibit concomitant high mTORC1 and AMPK activity suggesting that high mTORC1 signalling contributes to defective autophagy while AMPK may act to sustain residual autophagy (*58*). Thus, mTORC1 inhibitors or AMPK activators may be valuable treatment options for LRBA-deficient patients (*59, 60*). In fact, several clinical reports on patients with LRBA deficiency demonstrated that treatment with Sirolimus, an mTOR inhibitor, improved their clinical symptoms (*4, 61, 62*).

In conclusion, we found that LRBA facilitates autophagy through the interaction with PIK3R4 and FYCO1. Loss of these interactions resulted in low PI(3)P production, enlarged autophagosomes, aberrant trafficking and positioning of autophagosomes and lysosomes, leading to a reduced autophagosome-lysosome fusion, and thereby an inefficient degradation of cargo proteins *via* autophagy. Accumulation of cargo/antigenic peptides, particularly in the same vesicles that contain MHC-II molecules, results in enhanced antigen presentation leading to a stronger T-cell driven proinflammatory immune response in the context of LRBA deficiency.

## Material and Methods

### Study design

The objective of this study was to elucidate the molecular role of LRBA in autophagy and to assess the impact of LRBA loss on the regulation of the immune response, as LRBA-deficient patients suffer from a severe immune dysregulation phenotype. A combination of *in silico* and *in vitro* studies was used to identify two novel LRBA interactors, PIK3R4 and FYCO1. Both proteins are essential for autophagy, particularly for autophagosome membrane formation and autophagosome movement. Various *in vitro* and *ex vivo* assays were performed to explore how LRBA deficiency affects the different stages of autophagy including, PI(3)P production, recruitment of DFCP1 and WIPI2, size, maturation and movement of autophagosomes, autophagosome/lysosome fusion, as well as cargo degradation. In addition, we evaluated MHC-II-restricted antigen presentation, an autophagy-dependent process, and the subsequent T-cell cytokine release.

### Study protocol

Collection of PBMCs from healthy donors and LRBA-deficient patients to generate LCL cell lines was approved by the ethics committee of the University of Freiburg, Germany, vote n° 290/13.

### Generation of a LRBA-KO HEK293T cell line and a HaCat cell line by CRISPR-Cas9 and a LRBA knock-down HeLa cell line by shRNA (shLRBA cells)

LRBA-KO HEK293T and HaCat cells were generated using the Alt-R-CRISPR-Cas9 system according to the manufactureŕs instructions (www.idtdna.com). The gRNA CCACCAACAGGTGATGACGG specific for exon 2 of human LRBA, was inserted into the cells by electroporation together with the crRNA : tracrRNA complex and the Cas9. Following 48 h incubation, cells were single cell sorted by flow cytometry. LRBA-KO clones were validated by Western blot for LRBA expression abolishment and by Sanger sequencing for mutation identification (fig. S5, A-D). shLRBA HeLa cells were generated by lentiviral transfection with the vector pLKO.1 (SIGMA) containing the shRNA sequence: CCGGGCAGAAGATATTCACAGACATCTCGAGATGTCTGTGAATATCTT-CTGCTTTTTTG (TRCN0000148136, SIGMA) according to the manufactureŕs instructions (fig. S5E).

### Human lymphoblastoid B cell line generation

Lymphoblastoid B cell lines (LCL) were generated from B cells from three HD and two LRBA-deficient patients (P1: c.2004+2A>G; P2: p.S2713Hfs) after isolation from PBMCs by negative selection, and incubation 1:1 cell: EBV containing-media for four days. LCL actively proliferate, secrete antibodies and express LRBA (fig. S5F).

### Mice

*Lrba* knock-out mice (*Lrba^-/-^*) were kindly provided by Prof. Manfred Kilimann, Max-Planck-Institut für Experimentelle Medizin (MPIEM), Göttingen, Germany. They were generated on a C57BL/6N background by homologous recombination producing a loss-of-protein deletion of exon 4 of *Lrba* (*63*).

### Plasmids

Human LRBA was cloned in seven different fragments into pCIneo FLAG vectors (Promega). These plasmids were kindly provided by Dr. Bernice Lo from the National Institute of Health, Bethesda, USA (*6*). mCherry-GFP-LC3 plasmid was a kind gift from Dr. Ian Gentle from the University of Freiburg, Germany, whereas GFP-FYCO1 was kindly provided by Dr. Christian Behrends from the University of Frankfurt, Germany. Human LRBA full length tagged to Myc-DDK was purchased in Origene (RC17204). pVitro-hygro-N-myc-hVps34/Vps15-C-V5-his-plasmid (Addgene #24055), pEGFP-Atg14L (Addgene #21635) and pEGFO-C1-hUVRAG (Addgene # 24296) were provided by Jonathan Backer (*19*), Tamotsu Yoshimiri (*64*) and Noburo Mizushima (*65*), respectively. GFP-LC3 and MP1-LC3 plasmids were previously generated in the laboratory of Prof. Christian Münz from the University of Zürich, Switzerland.

### Reconstitution of LRBA

Rescue experiments were conducted in LRBA-KO HEK293T and HaCat cell lines transfected with 2 μg of WT LRBA plasmid tagged with myc (Origen) using Lipofectamine 2000 (Invitrogen). These reconstituted cells lines are called Myc-LRBA in this manuscript. Details of the transfection protocol are described in the “Transfections and Immunoprecipitations” section below.

### Proximity Ligation Assay

In situ PLA was performed using Duolink kit (Sigma Aldrich) in LCL cells from healthy donors (HD) 1 and 2, and Patient 1 and 2. Cell fixation and permeabilization was performed with 4% PFA and 0.1% Triton X-100, respectively. Incubation with primary antibodies was followed by incubation with secondary antibodies that are conjugated with oligonucleotides, PLA probe anti-mouse (or anti-goat) MINUS and PLA probe anti-rabbit PLUS. After ligation with DNA oligonucleotides and amplification with a DNA polymerase, the amplified product was detected as a fluorescent signal with a confocal microscope (Zeiss LSM700). Signal quantification was performed using Duolink Image Tool (Sigma Aldrich).

### Transfections

HEK293T cells were co-transfected with 2 μg of one of the seven Flag-tagged-LRBA-fragment plasmids or Myc-LRBA plasmid, plus 2 μg of Myc-PIK3R4, GFP-FYCO1, GFP-ATG14L, HA-VPS34 or GFP-UVRAG using Lipofectamine 2000 Turbofect (Invitrogen) according to the manufactureŕs protocol. Cells were harvested after 48 h and lysed with IP buffer (50 mM Tris-HCl pH 6.8, 150 mM NaCl, 0.2% NP-40, 1 mM EDTA), plus 1x complete EDTA-free protease inhibitor cocktail (Roche). Lysates were collected and used for immunoprecipitation experiments.

### Immunoprecipitations

LRBA IP, or flag-tagged LRBA IP or GFP IP were performed using 200 µg of cell lysates from HEK293T cells and 1 µg of either rabbit anti-LRBA, or mouse anti-FLAG antibody (Sigma Aldrich), mouse anti-PIK3R4 (Novus) or mouse anti-GFP (Santa Cruz). Then, 40 µl of Dynabeads protein G (Thermo Fischer Scientific) were added to the lysate/antibody mix and incubated overnight. Beads were washed with lysis buffer, and proteins were eluted with 2% SDS and resuspended in Laemmli buffer. 20 µl of the eluted proteins were separated by SDS-PAGE, blotted and detected by immunoblotting.

### Immunoblotting and antibodies

Cell lysates were generated with RIPA buffer (50 mM Tris, 1 % NP-40, 0.5 % sodium deoxycholate, 100 mM NaCl, 1 mM EDTA, 0.1% SDS) + 1x complete EDTA-free protease inhibitor cocktail (Roche). Total protein concentrations were determined by bicinchoninic acid Protein Assay (Thermo Fisher Scientific). Protein lysates were size-fractionated by SDS-PAGE and electro transferred to a PVDF membrane in a wet blotting system for 1.5 h at 45 V. After blocking with 5% milk in TBST (20 mM Tris, 150 mM NaCl, 0.1% Tween 20), the membranes were incubated at 4°C with any of the following primary antibodies: anti-AMPK total (Cell Signaling), anti-AMPK (pT172; Cell Signaling), anti-FLAG (Sigma Aldrich), anti-GFP (Santa Cruz), anti-LC3 (Novus), anti-LRBA (Sigma Aldrich), anti-Myc (Cell Signaling), anti-PIK3R4 (Novus), anti-p62 (Enzo Lifesciences), anti-S6K total (Cell Signaling), anti-S6K70 (pT389; Cell Signaling). After overnight incubation with the primary antibodies, membranes were immunodetected with their corresponding secondary HRP-coupled antibody (Santa Cruz). Membranes were washed and developed with Signal Fire or Signal Fire Plus chemiluminescent substrates (Cell Signaling). HRP-linked anti-Tubulin (Proteintech) and anti-GAPDH (Sigma) were used as a loading control. Peroxidase activity was detected with the Fusion SL device (Peqlab).

### Autophagy induction

Naïve B cells (CD43^-^B220^+^CD3^-^) were obtained by negative selection (mouse CD43 Ly-48 MicroBeads; MACS Miltenyi Biotec) from spleens of WT and *Lrba^-/-^* mice, followed by stimulation with LPS from *Escherichia coli* 055: B5 (20 µg/ml, Sigma-Aldrich) for 3 days in complete RPMI medium (L-glutamine, 10% FCS, 100 μg/mL streptomycin, 100 U/mL penicillin, 10 mM HEPES and 1 mM sodium pyruvate) with or without 100 nM of Bafilomycin A1 (InvivoGen) for 3 h. Next, cell pellets were either used for LC3-II detection by immunoblotting or for morphology and LRBA cellular localization analysis by electron microscopy. shControl and shLRBA HeLa cells were seeded at 0.3*10^6^ cells in 6 well-plates followed by treatment for 16 h or 24 h with 5 μM of MG132 (InvivoGen) alone or in combination with 100 nM of Bafilomycin A1 for 3 h.

### Measurement of Phosphatidyl inositol-3 phosphate (PI(*3*)*P*)

WT and LRBA-KO HEK293T cells were seeded and treated with 333 nM of Torin 1 (Invivogen) or with 1 nM of VPS34-IN (Biomol) for 4 h. After cell collection, cells lysis and extraction of neutral and acidic lipids, PI(3)P was obtained from the organic phase as described before (*66*). Finally, PI(3)P was measured using the PI(3)P Mass ELISA kit (Echelon) according to the manufactureŕs protocol.

### Fluorescence microscopy

To visualize autophagosomes, shWT and shLRBA HeLa cells were transfected with GFP-LC3 using Lipofectamine 2000 (Invitrogen). Lysosomes were visualized with LysoTracker Red (Invitrogen), according to the manufacturer’s instructions. To visualize autophagolysosomes, WT and LRBA-KO HEK293T cells were either stably transduced with the mCherry-GFP-LC3 tandem vector (kindly provided by Dr. Ian Gentle, University of Freiburg), or stained endogenous LC3B) and Lamp2 proteins. Cells were treated for 4h with 333 nM Torin 1 (Invivogen) alone or in combination with 100 nM Bafilomycin A1 (Invivogen). To visualize endogenous expression of DFCP1 and WIPI2 expression, WT and LRBA-KO HEK293T cells were treated for 1 h with 50 µM Rapamycin (Invivogen), 333 nM Torin 1 (Invivogen), 100 nM Wortmannin (Invivogen) or incubated in EBSS medium (Thermo Fisher) for starvation. After treatments, cells were fixed for 10 min with 4% PFA (Sigma Aldrich) and permeabilized with either 0.05% Saponin (Sigma Aldrich) or 0.1% Triton X-100. Endogenous LC3 and Lamp2 expression was detected after 1h incubation with anti-LC3B (Novus) and anti-Lamp2 (ThermoFisher) antibodies, followed by 45 min incubation with secondary antibodies anti-rabbit-AlexaFluor555 and anti-mouse-AlexaFluor647 (Cell Signaling). DFCP-1 and WIPI2 expression was detected after overnight incubation with anti-DFCP1 (LS-bio) or anti-WIPI2 (Abcam) antibodies, followed by 1h incubation with anti-mouse or anti-rabbit AF488 or AF647 antibodies (Cell Signaling). After staining, cover slides were mounted with DuoLink In Situ Mounting Medium with DAPI (Merck). The images were acquired using Zeiss LSM710 or LSM880 confocal microscope. The number of dots per cells and colocalization was assessed with Image J. The autophagosome size was assessed using Imaris 9.7 (Oxford Instruments). Autophagosomal size was determined from binary images using the analysis particle module of FIJI software (version X), with a particle size from 0 to 20 µm2. Autophagosomal mobility was evaluated by time lapse movies of autophagosomes acquired in individual frames color-coded and projected into one image, using the temporal color-coding module of the Zeiss Zen Black software. Autophagosomes were followed for 25 frames and area and velocity were quantified in pixels.

### LC3B co-localization with HLA-DR molecules in MIICs experiments

HaCat cells were seeded in glass cover slips placed inside 24-well plate and stimulated overnight with IFN-ɣ to induce MHC-II expression. Cells were treated 6 h with 20 µM of chloroquine, and then with 4% PFA for 15 min at RT in the dark. Permeabilization was performed with 0.1% Triton X-100 for 5 min at RT, followed by three PBS washes. Then cells were saturated with blocking buffer (1% FBS-PBS) for 1 h at RT. Primary antibodies were diluted in the blocking buffer and incubated with the cells for 1 h at RT. Primary antibodies used: rabbit anti-LC3B (MBL/Novus) and mouse anti-HLA-DR (Cell Signaling). Three PBS washes were performed before staining with Alexa Fluor 488/555-conjugated goat anti-mouse or anti-rabbit (Thermo Fischer) for 1 h at RT. After three PBS washes, cell nuclei were stained with DAPI for 5 min prior to mounting the cover slip onto a glass slide using DAKO mounting medium (Agilent). Cells were visualized through 63x, 1.4 NA oil immersion objective with a confocal laser scanning microscope (SP8 upright, Leica or LSM710, Zeiss) and images were processed with Fiji software.

### Membrane flotation assay

Cells from three 15 cm dishes were washed twice with PBS and harvested. The cell pellets were collected after centrifugation at 800 x g for 5 min and resuspended in homogenization buffer (250 mM sucrose, 20 mM HEPES-KOH pH 7.4, 1 mM EDTA and complete EDTA-free protease inhibitor cocktail (Roche)). Cells were lysed by 40 strokes in a glass Dounce homogenizer (WHEATON® Dounce Tissue Grinder). Unbroken cells and debris were removed by two centrifugation steps at 2,000 x g for 5 min. The supernatant was mixed with an equal volume of 50% OptiPrep (D1556-250ML; Sigma-Aldrich) in homogenization buffer. Discontinuous Optiprep gradients were prepared as described previously (*25*), in SW41 tubes (344059; Beckman Coulter) by overlaying the following Optiprep solutions in homogenization buffer: 2.4 ml of the diluted sample (25% Optiprep), 1.8 ml of 20% Optiprep, 2 ml of 15% Optiprep, 2 ml of 10% Optiprep, 2 ml of 5% Optiprep and 2 ml of homogenization buffer without Optiprep. The gradients were spin and 13 fractions of 0.95 ml were collected from the top and subjected to TCA precipitation. The final pellet was resuspended in sample buffer and incubated at 95°C for 5 min.

### Proteinase protection assay

Cells treated for 2 h with EBSS and 300 nM Torin 1, afterwards they were washed with PBS and collected by centrifugation at 500 x g for 5 min. Pellets were resuspended in homogenization buffer (250 mM sucrose, 20 mM Hepes-KOH pH 7.4, 1 mM EDTA and complete EDTA-free protease inhibitor cocktail) and lysed by 30 passages with a 25G needle. After two preclearing steps (2,000 x g, 5 min), cell membranes were pelleted by centrifugation at 20,000 x g for 30 min. The pellet was resuspended in 100 µl of homogenization buffer without EDTA or protease inhibitors, divided into three equal fractions and incubated in the presence or absence of proteinase K (100 µg per ml of sample) with or without 0.5% Triton X-100 for 30 min on ice. The samples were then subjected to TCA precipitation and resuspended in sample buffer.

### Pre-embedding immunoperoxidase and immunogold electron microscopy (EM)

Following fixation in 0.05% glutaraldehyde and 4% PFA in 0.1 M phosphate buffer (PB), cell pellets were embedded in agar and cut onto 100 µm slices using a Vibratome. Slices were blocked with 20% normal goat serum and incubated with anti-Lamp1 or anti-LC3 antibodies, followed by incubation with biotinylated anti-mouse and with goat anti-rabbit IgG coupled to 1.4 nm gold particles in 2% NGS/ 50 mM Tris-buffered saline. Slices were postfixed in 1% glutaraldehyde, and the gold particles were enlarged using the silver enhancement kit HQ-Silver from Nanoprobe. Slices were washed with ABC Elite Kit and peroxidase reaction was visualized by 3’3-diaminobenzidine. After washing, slices were postfixed with 0.5% OsO_4_/ 1% uranyl acetate, dehydrated in a graded series of ethanol, treated with propylene oxide and embedded in Durcupan resin. Slices were cut into 60 nm ultrathin sections, counterstained with lead citrate and viewed in a Zeiss LEO 906 transmission electron microscope. Images were taken using the sharp-eye 2k CCD camera and processed with ImageSP.

### Flow cytometry analysis

Intracellular expression of HLA-DR in IFN-γ-stimulated HaCat cells or intracellular expression of pS6k and pAMPK in LCL cells were determined as follows: 3*10^5^ cells were first resuspended in PBS and stained for fixable viability dye (eFluor780; Invitrogen) for 20 min at 4°C protected from light. Next, cells were fixed and permeabilized for 20 min using BD Cytofix/Cytoperm solution^TM^ (BD), and then washed twice with 1× Perm/Wash Buffer^TM^ (BD). Subsequently, cells were resuspended in 1× Perm/Wash Buffer and stained with conjugated anti-pS6K-Pacific Blue (Cell Signaling) and unconjugated anti-pAMPK (Cell signaling) for LCL cells, or with conjugated anti-HLA-DR (Invitrogen) for HaCat cells for 1 h at 4°C. After washing two times, a secondary antibody anti-rabbit IgG-FITC (Invitrogen) was added to LCL cells and incubated at 4°C for 25 min. Cells were then washed and acquired on a FACS Canto II (BD). Data analysis and calculation of the geometric mean fluorescence intensity (MFI) were performed using FlowJoTM 7.6.5 software (TreeStar Inc., Ashland, OR, USA). Gating strategy excluded doublets according to their FSC-H and FSC-A and dead cells, which were positive for fixable viability dye.

### In silico analysis

The LRBA protein interaction candidates were obtained from STRING and FUNCBase databases that provide known and predicted physical and functional protein-protein interactions. Only interactions with high confidence levels (>0.7) were selected from both databases for further validation. Network visualization was performed with the Cytoscape software provided for both databases.

### Statistical analysis

Statistical significance was calculated with a non-parametric two-tailed Mann–Whitney test, t-test or Two-Way-Anova using GraphPad Prism 6.0 software. A p-value of <0.05 was considered statistically significant (*p<0.05; **p<0.01; ***p<0.001; ****p<0.0001).

## Acknowledgements

We thank Dr. Bernice Lo (National Institute of Allergy and Infectious Diseases, Bethesda, USA), Dr. Christian Behrends (University of Frankfurt, Germany) and Dr. Ian Gentle (University of Freiburg, Germany) for kindly providing us with plasmids. We thank Dr. Michael Moutschen (University of Liege, Belgium) and Prof. Manfred Kilimann (University of Gottingen, Germany) for kindly sharing with us the shLRBA HeLa cell line and the *Lrba^-/-^* mice, respectively. In addition, we thank Jessica Rojas-Restrepo, Hanna Haberstroh, Türkan Güzel, Katja Malfertheiner, Vanessa Zeidler and Julika Neumann from our research group for their technical assistance during these past years, and to Dr. Virginia Andreani for her thorough review of this manuscript. We thank Dr. Marie Follo from the Light Core Facility Unit for her support and teaching of the microscopes. Schematic illustrations were created using Biorender licenses # QO24HLH0U1 and #KZ24HLGTRC.

## Author contributions

LG-D and BG designed the study. LG-D, LAL, CM, and CK designed the experiments. LG-D, ES, LAL, MCD, PS, SN, SR, PM and AR performed the experiments. LG-D, LAL, ES, MCD, PS, SG, SJ, MP, KT, ER, AR, CM and CK analyzed and interpreted the data. LG-D wrote the manuscript with contributions from all co-authors. LG-D, ES, MCD, LAL, PS, SN, SG, and AR prepared the figures. BG and LG-D secured funding for this work. All authors reviewed and approved the manuscript.

## Data availability

All data associated with this study are present in the paper or the Supplementary Materials. For original data, please contact

## List of Supplementary Materials

Materials and Methods

Fig. S1 to S5

Movie S1 to S2

References (*67–70*)

