## Supplementary Material for "LRBA balances antigen presentation and T-cell responses *via* autophagy by binding to PIK3R4 and FYCO1"

**Supplementary Material and Methods**

*Modelling of the complex of LRBA and PIK3R4/PIK3C3*

The structural modelling of the complex of LRBA and PIK3R4 was based on the finding that LRBA interacts *via* the WD40 domain with PIK3R4 (see results section). The 3D structure of the WD40 domain was modelled by homology modelling applying the structure of the homolog prokaryotic protein PkwA from *Thermomonospora curvata* (PDB-ID: 5YZV *(67)*). Sequence identity was 26% (39% shared similar residues). Since the first repeat sequence of the WD40-propeller of human LRBA is separated from the other WD40-repeats in sequence, this part of the domain was modelled based on a template of a single propeller and merged with the others, forming a full WD40 propeller domain. For the merging, a WD40 domain with an inserted propeller was used as a structural template (PDB: 3EWE; *(68)*). For the structural modelling of the complex of PIK3R4 with PIK3C3 the homologous structures of the Phosphatidylinositol 3-kinase VPS34 in complex with serine/threonine-protein kinase VPS15 from Saccharomyces cerevisiae was applied (PDB-ID: 5DFZ). Human PIK3C and the template share a sequence identity of 36% (with 54% similar residues). Human PIK3R4 and VPS15 share a sequence identity of 33% (50% similar residues). The modelling was performed with the homology modelling software Prime (Schrödinger Suite 2018-1, LLC). For the model of the WD40 domain of PIK3R4 we applied the modelling server WDSPdb 2.0 that accurately identifies WD40 repeats and includes them for the correct assignment of the beta sheets in the structural model *(69).* The WD40 domain was then connected with the kinase domain of PIK3R4 and finally energy minimized using the OPLS3 force field provided by the Schrödinger software package. The modelling of the complex of PIK3R4/PIK3C3 and LRBA was performed by applying the docking web server HADDOCK (High Ambiguity Driven Biomolecular DOCKing, version 2.2, *(70)*. As input constraints, the single exposed α-helix of the PIK3R4-WD40 domain (sequence motif: VGPSDD) and the smaller top surface of the WD40 domain of LRBA were defined as interacting regions. The top-ranked cluster with the highest negative Z-score value (-2.0) was selected for further analysis.

**Supplementary Figures**

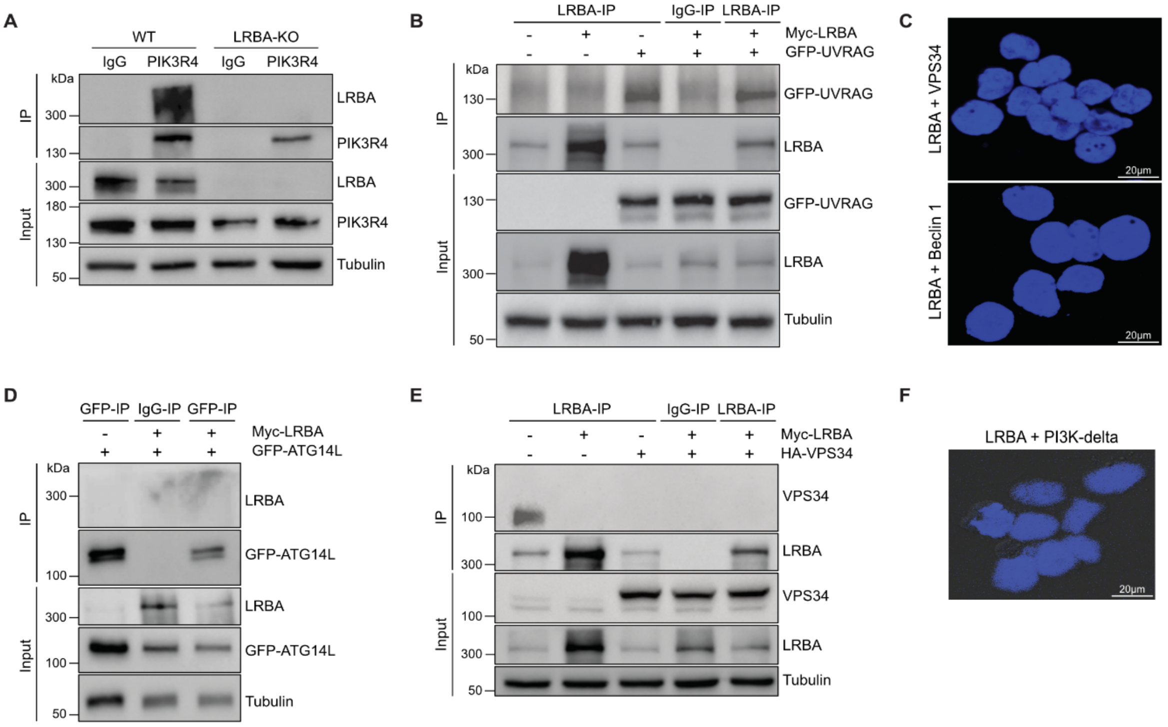


**Fig. S1**. **LRBA does not interact with other members of the PI3K-III complex core or with PI3Kdelta**. **(A)** Co-IP analysis of LRBA and PIK3R4 interaction at endogenous levels in WT and LRBA-KO HEK293T cells. Pull downs were performed with anti-PIK3R4 antibody and immunoblotted for LRBA. Tubulin expression was used as control. **(B)** HEK293T WT cells were transfected with Myc-LRBA and GFP-UVRAG plasmids. Pull downs were performed using anti-LRBA and immunoblotted for GFP. **(C)** Proximity ligation assay (PLA) showing the absence of close proximity of LRBA with VPS34 or Beclin-1 in LCL cells from a healthy donor under resting conditions. Nuclear DAPI staining is shown in blue. Scale bars, 20 µm. **(D)-(E)** HEK293T WT cells were transfected with Myc-LRBA plasmid and with either (D) GFP-ATG14L or HA-VPS34 (E). Pull downs were performed with anti-LRBA and immunoblotted with (D) anti-GFP or (E) anti-HA. **(F)** PLA showing the absence of proximity of LRBA with PI3Kdelta in LCL cells from a healthy donor under resting conditions. Nuclear DAPI staining is shown in blue. Scale bars, 20 µm.

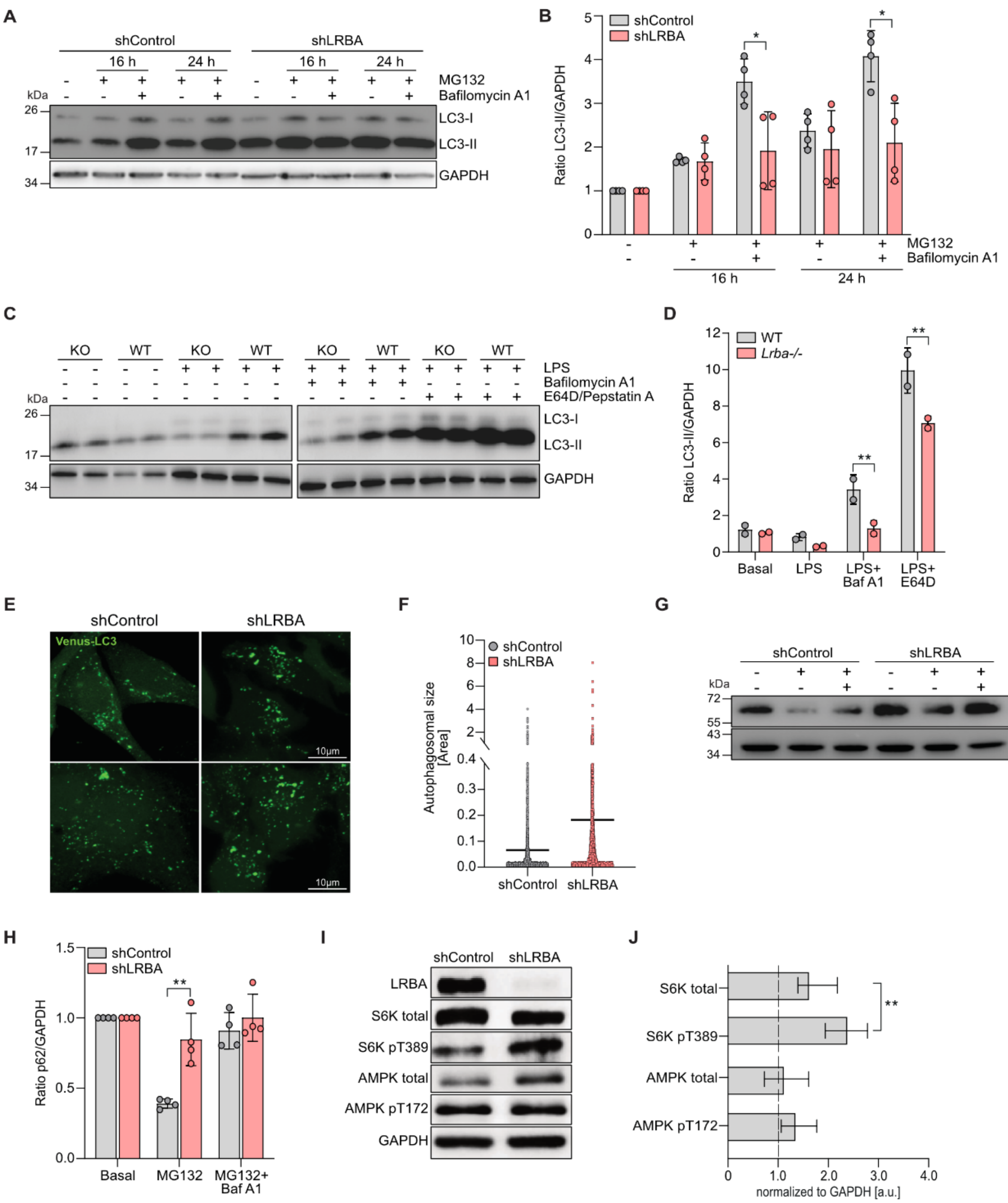


**Fig. S2. Abnormal autophagy flux in shLRBA Hela cells and B cells from *Lrba-/-* mice. (A)** Representative immunoblot analysis of the processing of endogenous unconjugated LC3-I to lipid-conjugated LC3-II in shControl and shLRBA HeLa cells at resting conditions or after 16 or 24 h of MG132 alone or in the presence of 100 nM Bafilomycin A1. **(B)** Densitometry analysis of LC3-II expression relative to GAPDH. **(C)-(D)** Isolated splenic murine naïve B cells from *Lrba-/-* and wild-type (WT) mice at day 0 or upon stimulation for 3 days with 20 µg/ml LPS alone or in the presence of 100 nM Bafilomycin A1 or protease inhibitors (1 mg/ml of E64D and Pepstatin A. **(C)**

Representative immunoblot analysis of LC3-I and LC3-II processing and Tubulin. **(D)** Densitometry analysis of LC3-II expression relative to Tubulin. **(E-F)** shControl and shLRBA HeLa cells were transfected with GFP-LC3 plasmid and incubated for 2 h with 100 nM of Bafilomycin A1 for microscopy evaluation. **(E)** Representative confocal microscopy images of LC3-transfected shControl and shLRBA HeLa cells (green), scale bar 10 µm. **(F)** Size of GFP-LC3 punctae in shControl (grey) and shLRBA (red) HeLa cells was determined from binary images using the analysis particle module of FIJI, with a particle size from 0 to 20 $\mu m^2$. **(G)** Representative immunoblot analysis of p62 and GAPDH in shControl and shLRBA HeLa cells at resting conditions or after 16 h of MG132 or in the presence of 100 nM Bafilomycin A1. **(H)** Densitometry analysis of p62 relative to GAPDH and normalized to basal conditions. **(I)** Representative immunoblot analysis of LRBA, S6K total, S6K pT389, AMPK total, AMPK pT172 and GAPDH performed in shControl and shLRBA-HeLa cells at basal conditions. **(J)** Bar graphs represent densitometry analysis of protein expression from HeLa cells using ImageJ. Quantifications of protein expression were previously normalized to GAPDH and to shControl (dotted line). Densitometry analyses were performed using ImageJ. The bars represent the means ± SD from n=4 (B) and (H), n=3 for (J) and n=2 (D) blots. Statistical analysis were performed using Mann-Whitney test, p-value *<0.05, **<0.01.

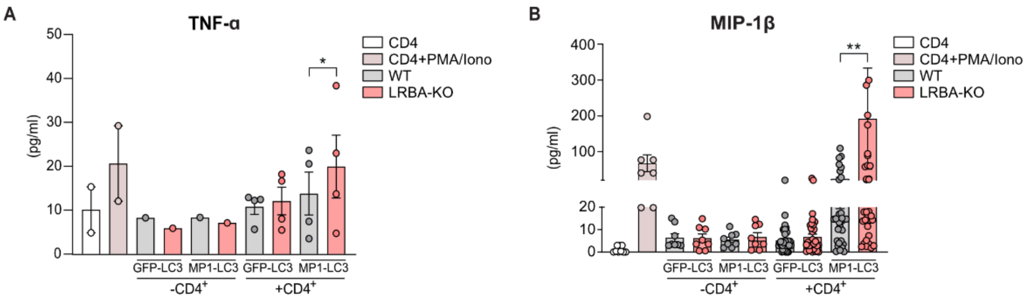


**Fig. S3. Loss of LRBA leads to increased TNF-α and MP-1β release upon antigen presentation. (A)-(B)** WT or LRBA-KO HaCat cells stably expressing GFP-LC3 or MP1-LC3 (target cells), and pre-treated with IFN-γ for 24h to up-regulate MHC-II molecules were cultured with a MP1-specific $CD4^+$ T-cell clone (effector cells) from a healthy donor for 20 h. Following incubation, TNF-α and MIP-1β were measured in the culture supernatants by ELISA. Bars represent the mean ± SD of **(A)** TNF-α (n=4) and **(B)** MIP-1β (n=4) secretion in WT (grey) and LRBA-KO (red) HaCat cells and each dot represents a replicate. Ratio paired student t-test, two-tailed, p-value *<0.05, **<0.0005.

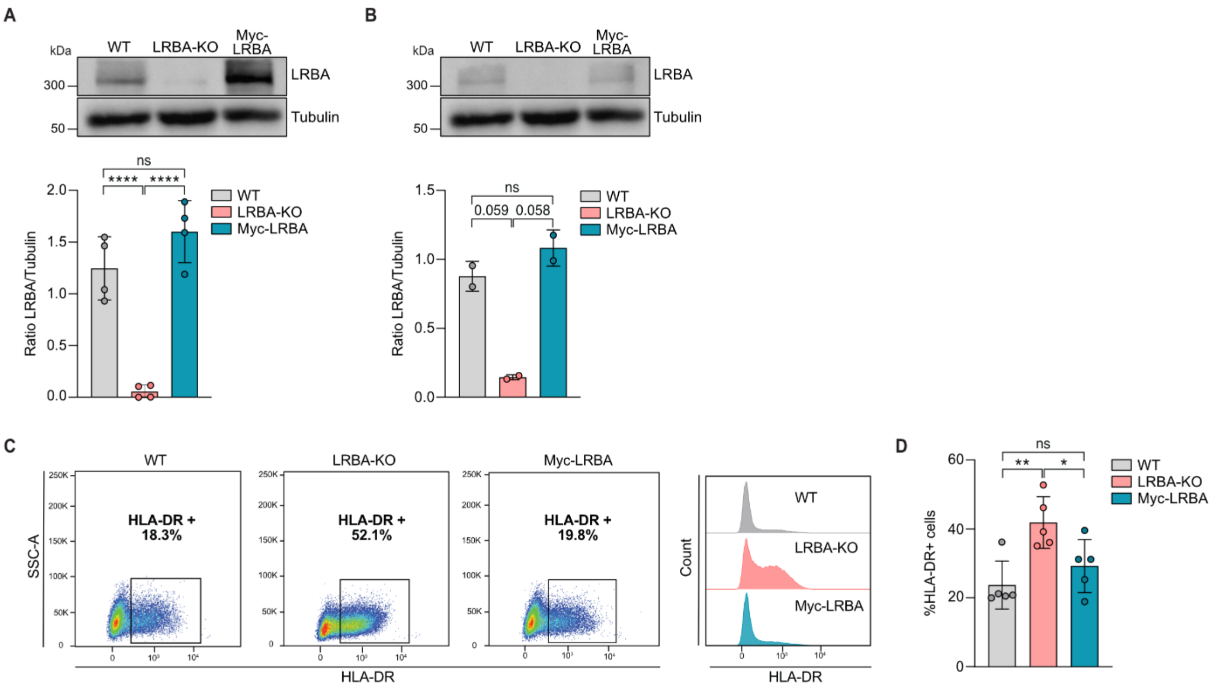


**Fig. S4. Ectopic reconstitution of LRBA expression restores normal levels of HLA-DR expression. (A)-(B)** Representative immunoblot analysis of LRBA expression in WT, LRBA-KO and Myc-LRBA reconstituted **(A)** HEK293T cells or **(B)** HaCat cells. Densitometry analysis of LRBA expression relative to Tubulin of WT (grey), LRBA-KO (red) and Myc-LRBA (teal). Bars represent the mean ± SD from (A) n=4 and (B) n=2 blots.**(C)-(D)** Flow cytometry analysis of HLA-DR expression at basal conditions and upon overnight stimulation with IFN-γ and 6 h Chloroquin stimulation are depicted in WT, LRBA-KO and Myc-LRBA HaCat cells. **(C)** Dot plots of HLA-DR positive cells and mean fluorescence intensity (MFI) histograms of HLA-DR expression. **(D)** Bars represent the quantification of WT (grey), LRBA-KO (red) and Myc-LRBA (teal) HaCat cells positive for HLA-DR from n=5 independent experiments. Welch's test was applied for (A), (B) and (D),p-value *<0.05, **<0.01, ***<0.001.

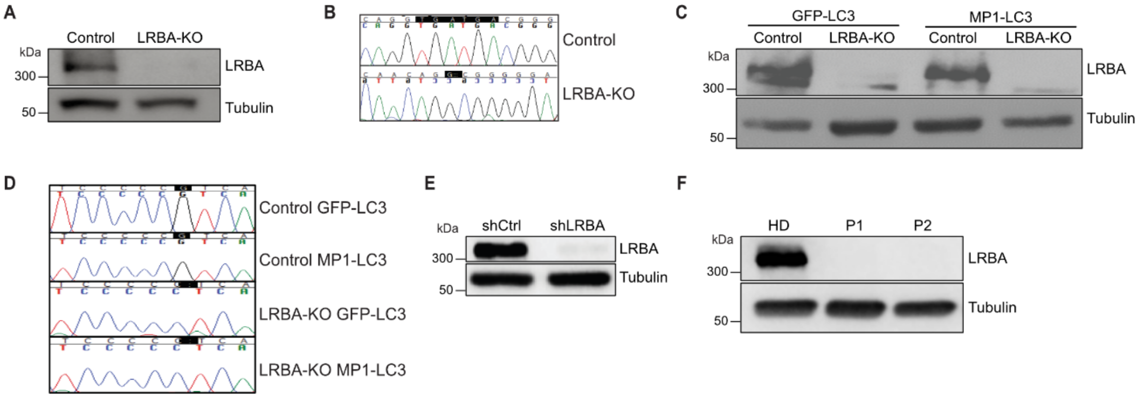


**Fig. S5**. **Generation of LRBA-deficient cell lines.** Immunoblot analysis of LRBA protein expression in WT and **(A)** LRBA-KO HEK293T or **(C)** LRBA-KO HaCaT cells. **(B)-(D)** sequencing analysis of *LRBA* exon 2 cells showing successful depletion of LRBA using the CrisprCas9 system in **(B)** HEK293T or **(D)** HaCaT cells. **(E)** Immunoblot analysis of LRBA protein expression in HeLa cells control and after treatment with shRNA for LRBA depletion (shLRBA). **(F)** Immunoblot analysis of LRBA protein expression in LCL cells from a HD and two LRBA-deficient patients.

**Supplementary Videos**

**Video 1 and Video 2.** WT and LRBA-KO cells were transfected with GFP-LC3 plasmid and incubated for 2 h with 100 nM Bafilomycin for live cell microscopy evaluation. Autophagosomes with increased size were found more frequently LRBA-KO cells (video 2) than in WT cells (video 1). In addition, autophagosomes with increased size presented slower movement in comparison to autophagosomes with regular size (video 2). Quantifications of these findings are depicted in Fig. 5, (A)-(C).
